# Identification of proteotoxic and proteoprotective bacteria that non-specifically affect proteins associated with neurodegenerative diseases

**DOI:** 10.1101/2023.10.24.563685

**Authors:** Alyssa C Walker, Rohan Bhargava, Michael Bucher, Amanda S Brust, Daniel M Czyż

## Abstract

Neurodegenerative protein conformational diseases (PCDs), such as Alzheimer’s, Parkinson’s, and Huntington’s, are a leading cause of death and disability worldwide and have no known cures or effective treatments. Emerging evidence suggests a role for the gut microbiota in the pathogenesis of neurodegenerative PCDs; however, the influence of specific bacteria on the culprit proteins associated with each of these diseases remains elusive, primarily due to the complexity of the microbiota. In the present study, we employed a single-strain screening approach to identify human bacterial isolates that enhance or suppress the aggregation of culprit proteins and the associated toxicity in *Caenorhabditis elegans* expressing Aβ_1-42_, α-synuclein, and polyglutamine tracts. Here, we reveal the first comprehensive analysis of the human microbiome for its effect on proteins associated with neurodegenerative diseases. Our results suggest that bacteria affect the aggregation of metastable proteins by modulating host proteostasis rather than selectively targeting specific disease-associated proteins. These results reveal bacteria that potentially influence the pathogenesis of PCDs and open new promising prevention and treatment opportunities by altering the abundance of beneficial and detrimental microbes.

## Introduction

Neurodegenerative protein conformational diseases (PCDs) are characterized by disturbances in proteostasis that result in the aggregation of disease-associated proteins, ultimately leading to tissue death.^1^ Alzheimer’s disease (AD), Parkinson’s disease (PD), Huntington’s disease (HD), and amyotrophic lateral sclerosis (ALS) are among the most prevalent neurodegenerative diseases, with AD recognized by the World Health Organization (WHO) as one of the leading causes of death worldwide.^2^ However, despite their increasing prevalence, the etiology and potential therapeutic targets remain obscure. The sporadic onset and variable severity of neurodegenerative diseases, along with their idiopathic nature, suggest the possibility of a triggering factor for their onset and progression. Multiple factors have been associated with the pathogenicity of PCDs, including an expanding body of evidence that suggests the involvement of microbes, but primarily those within the human gut microbiota (HGM). The HGM is considered an “organ” due to its production of essential proteins and metabolites, including vitamins, hormones, and neurotransmitters.^3,4^ Hence, dysbiosis of the gut microbiota has been linked to various ailments, including neurodegenerative diseases.^5^

The complexity of the microbiome has made it challenging to determine the precise role of bacteria in the pathogenesis of neurodegenerative diseases. In addition, most of the evidence that associates bacteria with the occurrence of neurodegenerative diseases is based on correlation.^6^ Interestingly, correlational evidence does not demonstrate any selectivity between different neurodegenerative diseases, despite each disease featuring a unique, specific culprit protein species. For example, a lower abundance of *Prevotella* spp. has been observed in patients with different PCDs, including PD, and ALS.^7–15^ Due to this lack of specificity, we hypothesized that bacteria could be affecting these diseases through the host proteostasis network— upstream of protein aggregate formation. This hypothesis is further supported by our previous study in which we identified gram-negative enteropathogens that significantly disrupt proteostasis across tissues in *C. elegans.*^16^ The utility of *C. elegans* as a model to study host-microbe interaction is strengthened by its unique ability to be colonized by a single bacterial strain. Such a feature simplifies the complexity of the microbiome, allowing us to study the effect of individual species on host proteostasis.

Here, we characterized the effect of 229 unique bacterial isolates from the Human Microbiome Project on the aggregation of disease-associated proteins in *C. elegans*, using transgenic nematodes expressing Aβ_1-42_, α-synuclein, and polyglutamine (PolyQ). Surprisingly, our results suggest that bacteria-mediated enhancement or suppression of host protein aggregation is not specific to any particular culprit protein species. Instead, our results demonstrate that bacteria broadly affect the aggregation of metastable proteins present within the host proteome. Furthermore, we also observed that the proteostasis-modulating effects of intestinal bacteria reach distal tissues. Thus, our results indicate that bacteria influence host proteostasis, ultimately affecting the ability of both proximal and distal tissues to buffer protein folding. To date, our results provide the most comprehensive characterization of the effect of individual constituents of the human microbiome on host proteostasis. These results reveal the bacterial contribution to the pathogenesis of AD, PD, HD, ALS, and possibly other PCDs. Collectively, our results provide a framework for the development of microbiome-based risk factor assessments and disease management strategies.

## Results

### Characterization of the human microbiome on host proteostasis

We obtained a comprehensive Human Microbiome Project collection of 229 unique bacterial isolates from BEI Resources and assessed their effect on *C. elegans* proteostasis. The method used to conduct this experiment is illustrated in Figure 1. The collection encompasses isolates from a range of diverse phyla and a variety of anatomical sites (Figure 2). In our previous study, we used aggregation-prone polyQ tracts as a sensor of the protein folding environment to demonstrate that gram-negative pathogens disrupt host proteostasis.^16^ Among animals carrying intestine-, muscle-, and neuron-specific polyQ::YFP, we previously found that proteostasis in the intestine was most robustly affected by the colonizing bacteria.^16^ As such, we performed our initial screen using the intestinal polyQ model (Figure 2). To eliminate the possibility that bacteria will affect *C. elegans* development, worms were grown to adulthood prior to being transferred to the bacterial strains of interest. polyQ aggregation was assessed by fluorescent microscopy. To ensure the validity of our results, we also assessed polyQ aggregation by western blotting. Validating our approach, both quantitative methods yielded similar results.

**Figure 1.**
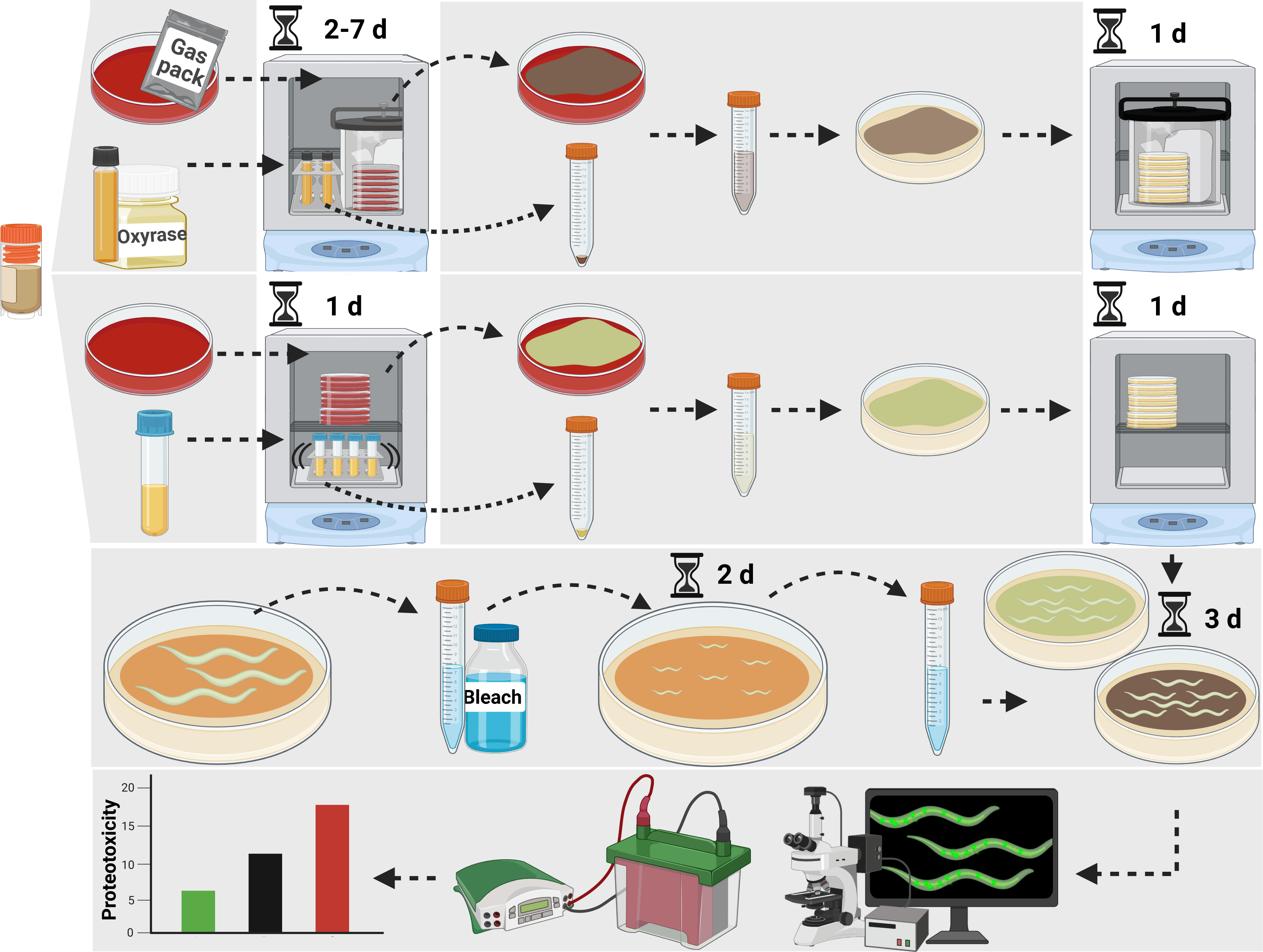
Schematic illustrating the method used to assess the effect of bacterial isolates on *C. elegans* proteostasis. Details can be found in Star Methods.

**Figure 2.**
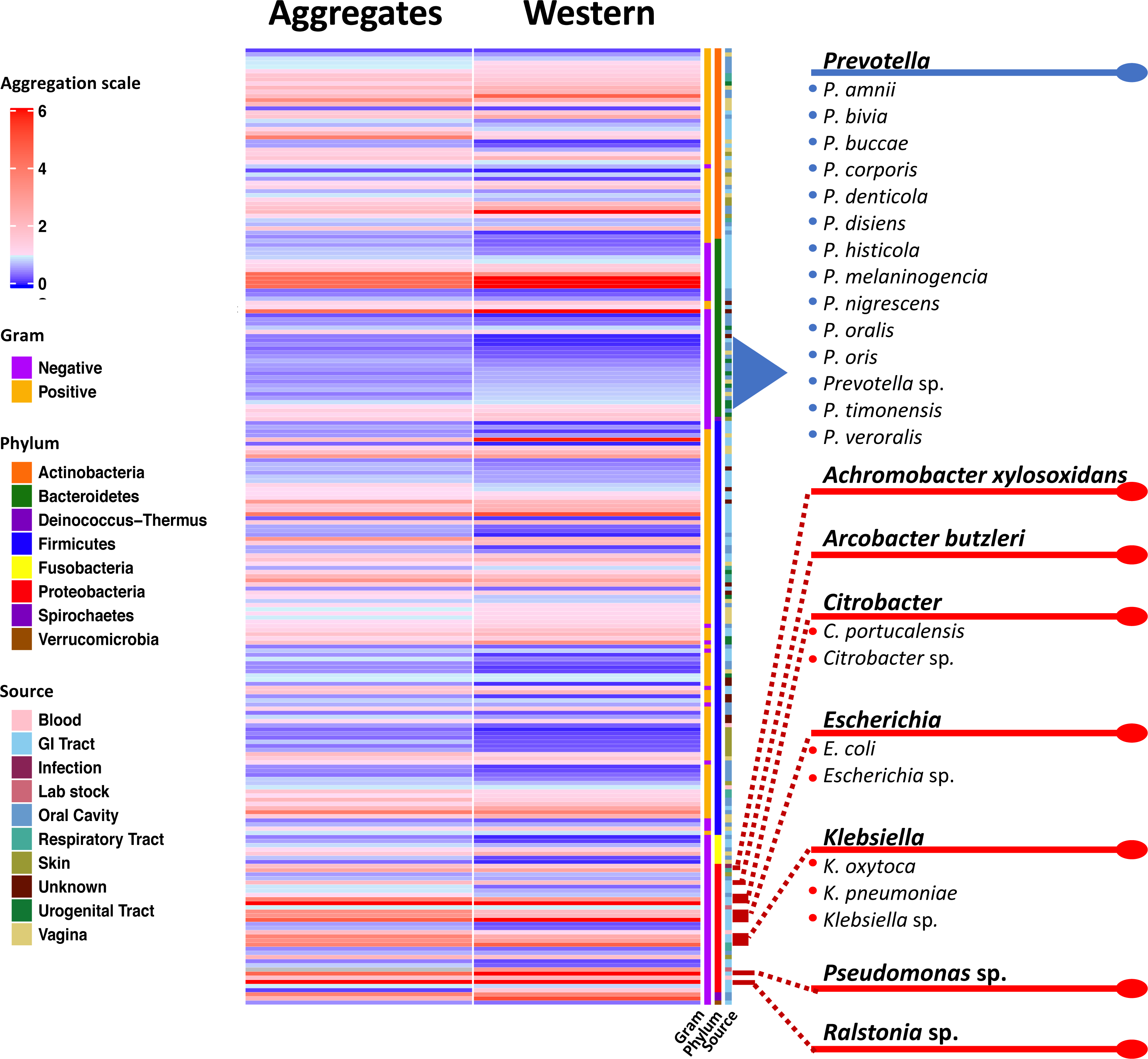
Heat map showing the extent of the screen that identified the effect of human bacterial isolates on *C. elegans* intestinal polyQ aggregation. Aggregation data are normalized to worms fed control *E. coli* OP50.

Bacteria from the isolate collection exhibited a differential effect on *C. elegans* proteostasis as indicated by increases and decreases in polyQ aggregation compared to worms fed control *E. coli* OP50 (Figure 2). Figure 2 summarizes the screen revealing bacteria that most robustly affected host proteostasis. For example, *Prevotella* was the only genus that consistently resulted in low polyQ aggregation across all 229 strains tested (Figure 2). Conversely, significant host proteotoxicity was observed in nematodes that were fed *Achromobacter xylosoxidans* and *Arcobacter butzleri,* as well as *Citrobacter, Escherichia, Klebsiella, Pseudomonas,* and *Ralstonia* spp. (Figure 2). A detailed list of all bacterial strains and their effect on polyQ aggregation is summarized in Table S1. Intriguingly, our data align with previous research linking the depletion or enrichment of these bacteria in patients with PCDs, which is further described in the discussion section of the present study. To our knowledge, this is the first-ever comprehensive screen that assessed the effect of human bacterial isolates on host proteostasis.

### *Prevotella* spp. mitigate the proteotoxic aggregation of diverse culprit proteins

Out of the 229 strains tested, the *Prevotella* genus had the highest number of members that suppressed polyQ aggregation. We further retested 17 *Prevotella* spp. using the intestinal polyQ44 model. To increase the robustness of the response, we started feeding animals test bacteria immediately after hatching. Although we observed an enhanced suppression of aggregation for most retested strains, there were few isolates that did not significantly inhibit polyQ aggregation, likely due to the timing of feeding. Furthermore, as expected, feeding worms test bacteria beginning at the L1 larval stage affected development (Figure 3A, “Aggregates”). Out of all strains, *P. buccae, P. oris*, and *P. corporis* significantly suppressed polyQ aggregation without causing any detectable developmental delay (Figure 3A, “Aggregates”). Therefore, we followed up with these three strains. Motility defects are associated with neurodegenerative PCDs. As such, we assessed whether the three *Prevotella* spp. alleviate aggregate-dependent motility defects caused by culprit proteins expressed in the intestine and distal tissues. We used our well-established and validated time-off-pick (TOP) assay that relies on *C. elegans* motility as a readout of proteotoxicity.^16,17^ Consistent with their suppression of polyQ aggregation, all three strains alleviated aggregate-dependent motility defects with *P. corporis* having the strongest effect (Figure 3A, “Motility”). To determine whether the observed suppression is dependent on polyQ, we used N2 wild-type worms and a model expressing a shorter polyglutamine tract, polyQ33. The results revealed no effect on the N2 worms and a less robust suppression of the motility defect for polyQ33, indicating that *Prevotella* suppressed proteotoxicity.

**Figure 3.**
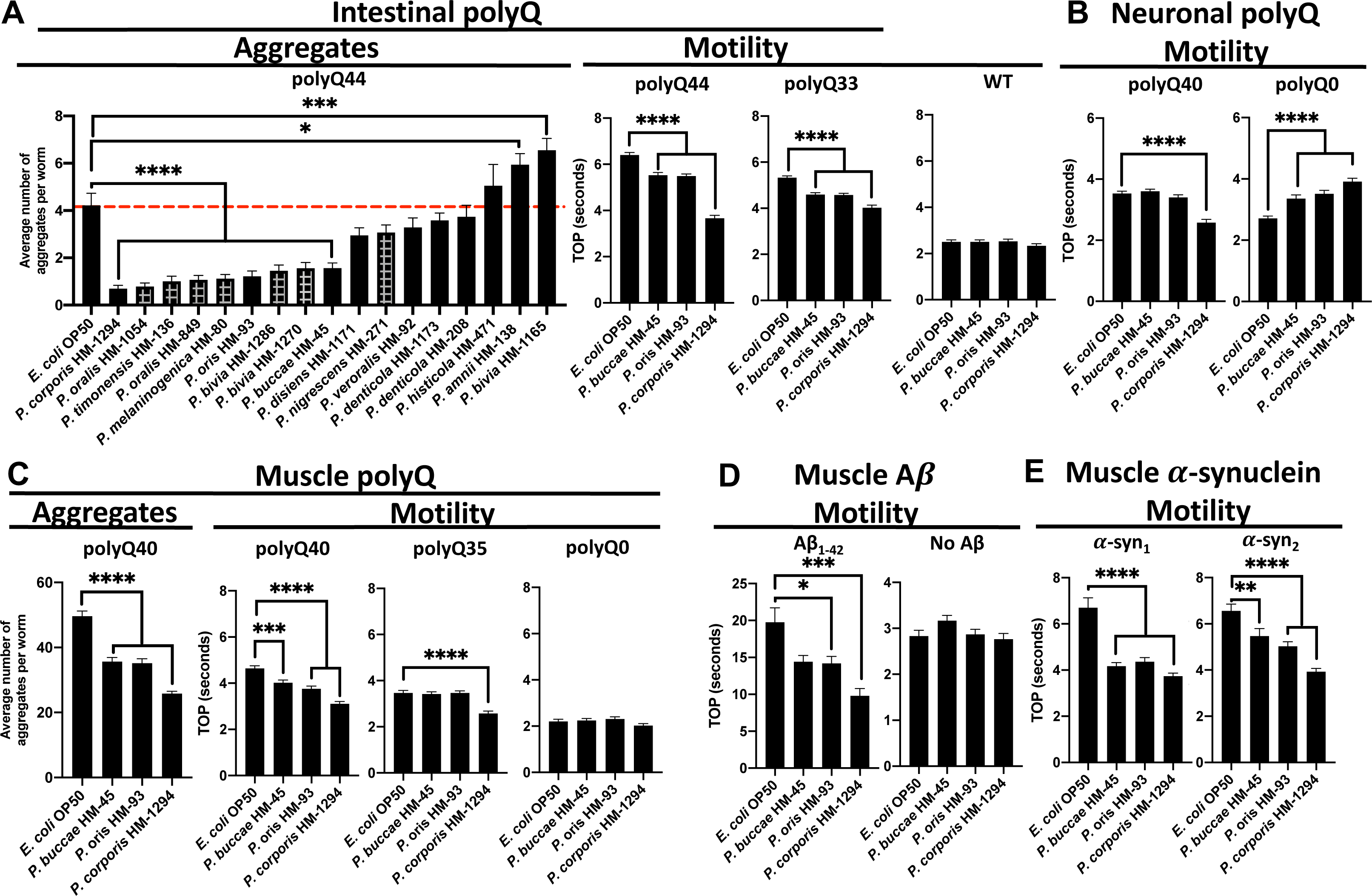
The effect of *Prevotella* spp. on proteins associated with neurodegenerative diseases. A) The effect of *Prevotella* spp. on *C. elegans* intestinal polyQ aggregation (“Aggregates”) and the associated toxicity (“Motility”). Checkered bars represent bacteria-associated developmental delay. The three strongest suppressors of polyQ aggregation that did not affect development were tested using intestinal polyQ, B) neuronal polyQ, C) muscle polyQ, D) muscle Aβ_1-42_, and E) muscle α-synuclein. Data are represented as the average number of aggregates or TOP (seconds) per worm obtained from at least two independent experiments for a total of 30-60 worms. Error bars represent standard error of the mean (SEM). Statistical significance was calculated using one-way ANOVA followed by multiple comparison Dunnett’s post-hoc test (*p<0.05, **p<0.01, ***p<0.001, ****p<0.0001).

To determine whether *Prevotella* also affects polyQ in distal tissues, we employed the neuronal and muscle models. Because worms expressing the neuronal polyQ do not exhibit quantifiable aggregates, we used the TOP assay as a motility readout. Our results show that of the three strains, only *P. corporis* showed significant suppression of the motility defect in worms expressing neuronal polyQ40 (Figure 3B). Unexpectedly, *P. buccae, P. oris*, and *P. corporis* increased motility defects in the neuronal empty vector control (polyQ0); the reason for this is unknown, but importantly does not diminish the observed beneficial effects of *Prevotella* on neuronal polyQ. In a manner similar to the intestinal polyQ, we observed Prevotella-mediated suppression of aggregation and proteotoxicity in worms expressing muscle-specific polyQ (Figure 3C).

The association between the low abundance of gut *Prevotella* and diverse neurodegenerative PCDs suggests that the bacterial influence on these diseases is independent of the culprit protein species.^8–14^. As such, we hypothesized that bacteria might affect protein aggregation by modulating host proteostasis, and if this is the case, *Prevotella* spp. should enhance proteostasis in worms expressing various aggregation-prone proteins. To test the effect of Prevotella on other disease-associated proteins, we chose three muscle-specific models: one expressing Aβ_1-42_, and two expressing α-synuclein.^18–20^ These culprit proteins model Alzheimer’s and Parkinson’s diseases, respectively. Additionally, these models exhibit age-dependent proteotoxicity, which is consistent with the age-dependent progression of neurodegenerative diseases.^16,21,22^ To investigate progressive proteotoxicity in the Aβ_1-42_ and α-synuclein models,^18–20^ we assessed their motility between days three to five post-hatching (Figure S2). Nematodes expressing Aβ_1-42_ and α-synuclein had age-dependent motility defects that were markedly greater than those of the controls (Figure S2). We examined the effect of *P. buccae, P. oris*, and *P. corporis* on proteotoxicity associated with each of these three models. In a manner similar to the polyQ models, *P. buccae, P. oris*, and *P. corporis* reduced aggregate-dependent toxicity compared to wild-type control (Figure 3A), particularly with *P. corporis* having the strongest effect (Figure 3D, Figure 3E). Bacteria that strongly enhance or suppress host proteostasis induce notable differences in the aggregation of Aβ_1-42_, which parallels the extent of the motility defect (Figure S1). Both Aβ_1-42_ aggregation and the associated toxicity were suppressed by *P. corporis* and enhanced by *Pseudomonas aeruginosa* (Figure S1), a bacterium that strongly disrupted host proteostasis in our previous studies.^16,23^ All of our transgenic worms express exogenous proteins. To determine whether *Prevotella* can also provide protection against misfolding of endogenous proteins, we used a strain that carries a temperature-sensitive mutation in myosin heavy chain (UNC-54) that leads to its misfolding and paralysis at the restrictive temperature (25°C).^24,25^ In agreement with the proteoprotective effect against misfolding and aggregation of exogenous proteins, *Prevotella* suppressed the temperature-dependent motility defect at the restrictive temperature, further supporting its beneficial role (Figure S3). Collectively, our findings suggest the broader potential of *Prevotella* in mitigating the pathogenesis of neurodegenerative diseases.

### Proteotoxic bacteria enhance the aggregation and toxicity of PCD-associated proteins

To confirm the effect of the most robust proteotoxic strains identified in our original screen (Figure 2, Table S1), we assessed polyQ44 aggregation in animals fed gram-negative aerobes (Figure 4A, “Aggregates”). We concentrated on these specific bacteria because they did not affect worm development, which is known to influence proteostasis.^26^ We found that *Ralstonia* sp., *Achromobacter xylosoxidans*, *Pseudomonas* sp., and *K. pneumoniae* were the strongest inducers of polyQ aggregation (Figure 4A). As we have already demonstrated the proteotoxic potential of *P. aeruginosa* in our previous work,^16,23^ we focused on the three remaining species in follow-up experiments. To assess the proteotoxic effect of these bacteria, we employed the TOP assay to measure the motility of worms expressing aggregating polyQ44 and non-aggregating polyQ33. While we detected a significant enhancement in the motility defect in animals expressing polyQ44, no significant changes were observed in wild-type or in animals expressing polyQ33 (Figure 4A). These results indicate that the impairment of motility induced by bacteria is contingent on polyQ aggregation rather than general bacterial pathogenicity. The aggregate-dependent motility defects were also observed in worms expressing neuronal and muscle-specific polyQ; although, the phenotype was less robust as we did detect some decrease in motility in control animals (Figures 4B and 4C). *K. pneumoniae*, *Ralstonia* sp., and *A. xylosoxidans* also elevated muscle-specific polyQ aggregation (Figure 4C). To test the microbial influence on other disease-associated proteins, we employed worms expressing muscle-specific Aβ_1-42_ and α-synuclein. In agreement with the intestinal polyQ model, all of these bacterial isolates also induced proteotoxicity associated with these additional culprit proteins (Figures 4D and E). These results demonstrate that bacteria can enhance the proteotoxicity of diverse disease-associated proteins, supporting their role in the pathogenesis of neurodegenerative PCDs.

**Figure 4.**
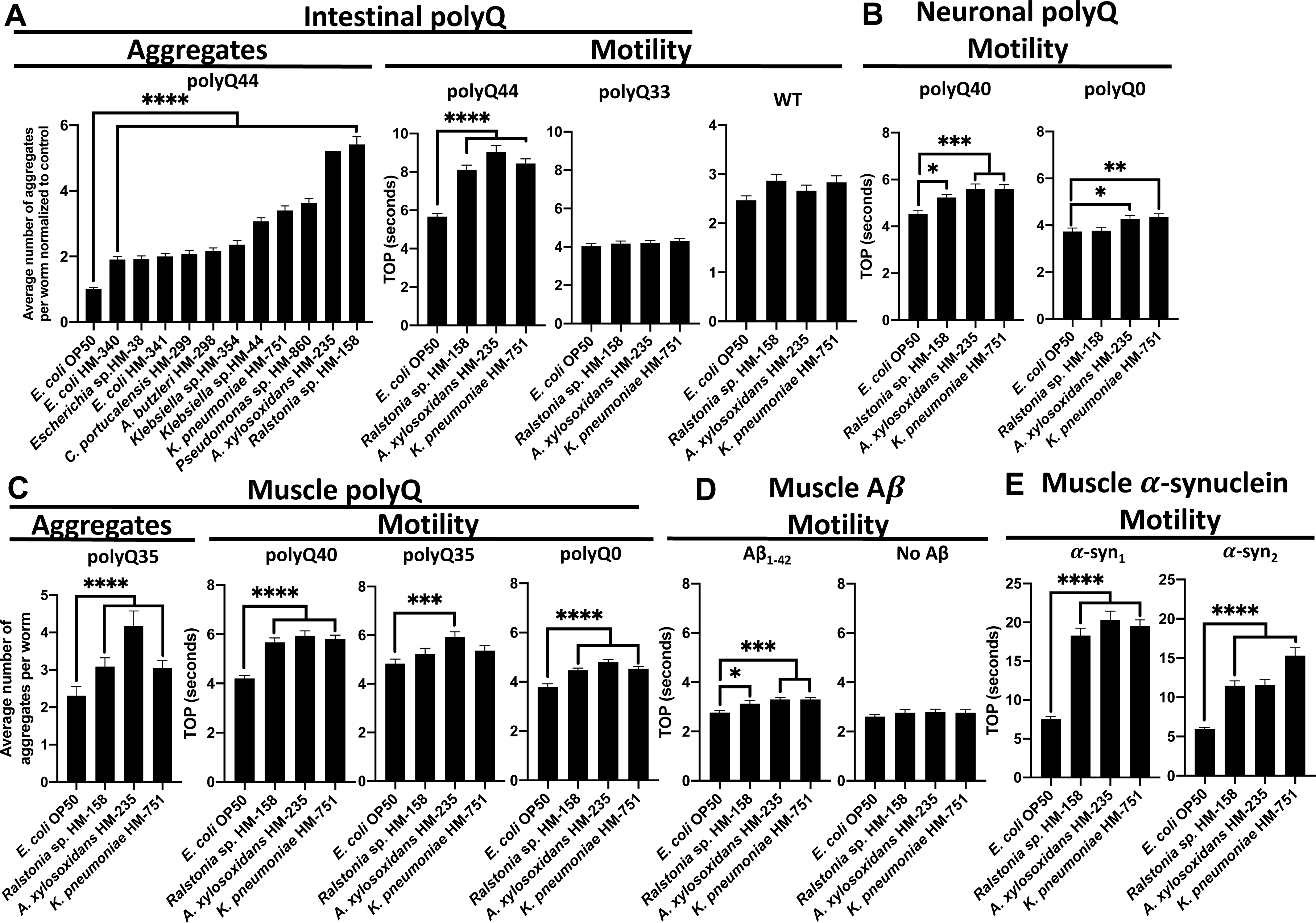
The effect of proteotoxic bacteria on proteins associated with neurodegenerative diseases. A) The effect of gram-negative, aerobic bacteria on *C. elegans* intestinal polyQ aggregation (“Aggregates”) and the associated toxicity (“Motility”). Three robust enhancers of polyQ aggregation were tested using intestinal polyQ, B) neuronal polyQ, C) muscle polyQ, D) muscle Aβ_1-42_, and E) muscle α-synuclein. Data are represented as the average number of aggregates or TOP (seconds) per worm obtained from at least two independent experiments for a total of 30-60 worms. Error bars represent SEM. Statistical significance was calculated using one-way ANOVA followed by multiple comparison Dunnett’s post-hoc test (*p<0.05, **p<0.01, ***p<0.001, ****p<0.0001).

## Discussion

In the present study, we screened a comprehensive collection of 229 unique bacterial isolates from the Human Microbiome Project for their ability to affect host proteostasis from within the intestinal milieu. Our follow-up experiments on the most robust beneficial and detrimental bacteria confirmed their influence on proteostasis across host tissues, affecting the stability of proteins associated with Alzheimer’s, Parkinson’s, and Huntington’s diseases. The phylogenetic analysis revealed clustering of select proteotoxic and proteoprotective bacteria, indicating that genetically related bacteria affect host proteostasis in a similar manner, but with a different magnitude (Figure S4). To our knowledge, this is the first comprehensive characterization of bacteria from human microbiomes on host proteostasis. Surprisingly, our data suggest that bacteria do not selectively target any specific host proteins associated with PCDs, but rather affect proteostasis in general, leading to the aggregation and proteotoxicity of any destabilized proteins present within the proteome, such as exogenous polyQ, Aβ_1-42_, and α-synuclein, tested in our experiments. While this is a generalized mechanism, there could be microbes that exclusively affect a specific disease.

Numerous studies that employed genomic analyses of microbial compositions in affected patients revealed a connection between comparable gut dysbioses and diverse neurodegenerative PCDs.^6^ These correlational studies from human subjects support our conclusion that bacteria impact protein stability by modulating host proteostasis. This mechanism is likely mediated by the interplay between host proteostasis and immune responses to bacteria.^27,28^

Many of the studies aiming to identify neurodegenerative PCD etiology or treat the disease have centered on host-targeted therapeutics, primarily focused on targeting the aggregating proteins or the affected cell types. However, this approach has not been successful in pre-clinical or clinical trials.^29,30^ Perhaps the focal point of the host-targeted approach occurs too late in the aggregation cascade given that the changes in the gut microbiota can happen prior to any clinical manifestation.^31,32^ Indeed, individuals with neurodegenerative PCDs have insufficient proteostatic capacity.^33^ While some studies have attempted to treat neurodegenerative PCDs by enhancing components of the host proteostasis network, these have not been successful.^29^ In support of our results indicating that bacteria affect host proteostasis, a shift towards microbial-centered therapeutic approaches has shown more promise in lessening disease symptoms. For example, eradication of *H. pylori* in AD patients was associated with improved disease presentation in a clinical trial and a population-based study.^34,35^ Interestingly, the beneficial effect of antibiotics on the progression of PCDs, when administered post-onset, is absent when antibiotics were used prior to disease onset; notably, general antibiotic use has been associated with increased risk for PCDs as well as gut dysbiosis.^36–38^ While *H. pylori* is a good clinical example of microbial contribution to neurodegenerative diseases, we did not expect to detect proteotoxicity associated with this bacterium in our *C. elegans* model due to a limitation of our approach that includes transferring and incubating bacteria at sub-optimal temperature – a factor that facilitates *H. pylori* virulence.^39^ Additional reports suggest that *H. pylori* can indirectly affect the composition of the human gut microbiota.^40^ Microbe-targeted therapeutic approaches hold promise and are further bolstered by a recent study that demonstrated that gut microbiota and serum derived from AD patients accelerate disease symptoms and affect neurogenesis in healthy rats and tissue culture, respectively.^41^

In our screen (Figure 2), *Prevotella* stood out as the only genus devoid of any species that exhibit proteotoxicity towards the host. Furthermore, our findings indicate that *Prevotella* spp. have a broad and suppressive effect on host protein aggregation, regardless of the specific disease-associated protein (Figure 3). Sequencing data suggest that a low abundance of *Prevotella* in the guts of patients with PCDs enhances disease pathogenesis.^7–14^ However, despite the fact that the *Prevotella* genus contains over 50 species, many studies report their results at the genus level.^42^ Our data indicate that various *Prevotella* species exhibit a differential effect on host proteostasis (Figure 3A). Such various effects of individual species could be explained by large variability in their genomes.^42^ The species, *P. intermedia*, *P. nigrescense*, and *P. melanigencia* have been consistently associated with infection,^43–45^ inflammation,^46–49^ and have also been linked to AD and associated mortality.^50^ Conversely, it has been shown that *P. buccae* and *P. corporis*, two strains of *Prevotella* that exhibited strong suppression of protein aggregation and the associated toxicity, were found not to induce inflammation.^51^ Instead, *P. buccae* and *P. corporis* were shown to induce the expression of mucin-associated membrane proteins MUC3 and MUC4.^51^ Interestingly, induction of MUC3 expression by a probiotic cocktail prevented adherence of enteropathogenic *E. coli.*^52^ MUC4 has been shown to be essential for maintaining mucus barrier function and intestinal homeostasis in mice, as its absence resulted in severe large intestinal bleeding and significant down regulation of antimicrobial peptides.^53^ Though preliminary, the evidence suggesting the protective role of *Prevotella* in maintaining intestinal integrity is interesting, as intestinal integrity is often compromised in people with neurodegenerative PCDs.^54^ Damage to the intestinal epithelium can lead to translocation of bacteria and bacterial products, resulting in systemic inflammation and the breakdown of the blood-brain barrier, contributing to a variety of diseases, including PCDs.^55,56^ The contrasting associations of different *Prevotella* species in host health and disease, highlight the importance of studying bacteria at the species level, as demonstrated in the present study, to unravel the precise roles of bacteria in disease pathogenesis.

The physiological relevance of our findings is supported by studies that focus on pathogenic bacteria and subsequently establish a connection with PCDs. For example, the robust induction of proteotoxicity by *K. pneumoniae* across all *C. elegans* models expressing different aggregation-prone protein species (Figure 4), is in agreement with a previous study that identified an overabundance of *K. pneumoniae* in the gut microbiota of individuals with AD.^57^ Furthermore, a positive correlation has been demonstrated between the presence of *K. pneumoniae* in the gut and elevated levels of the AD biomarker, C-reactive protein, in blood samples of AD patients.^58^ Another study found an elevated abundance of *Ralstonia* in mucosal samples from individuals with PD,^59^ which is another bacterium that was robustly proteotoxic to our *C. elegans* models (Figure 4). Interestingly, this bacterium was also found to be more prevalent in individuals with autism,^60^ which is another disease that has been associated with the presence of misfolded proteins.^61–64^ Autism has also been associated with both *Achromobacter* and *Pseudomonas* genera.^65,66^ Both *Achromobacter* and *Pseudomonas* are linked to cystic fibrosis, which is another PCD that does not feature neurodegeneration;^67,68^ *Achromobacter* was reported to exacerbate the disease,^69^ and *P. aeruginosa* is a predominant pathogenic species that colonizes the lungs of CF patients.^70^ *A. xylosoxidans* induced proteotoxicity in all our disease models (Figure 4), along with *Pseudomonas*, which displayed a notable increase in polyQ aggregation (Figure 4A, “Aggregates”). Furthermore, we previously demonstrated that *P. aeruginosa* exert robust proteotoxic effects on *C. elegans.*^16,23^ The presence of *Pseudomonas,* in conjunction with another bacterial genus, was successfully used to differentiate individuals with AD from control subjects.^71^ Collectively, the aforementioned studies are in agreement with our results and further support the wide-ranging impact of bacteria on protein folding diseases. It is remarkable that the abovementioned bacteria induce proteotoxicity consistently across all our *C. elegans* models expressing distinct disease proteins. The convergence of our results with existing literature linking these bacteria to PCDs strongly reinforces the notion that bacteria affect the host proteostasis network, broadly affecting protein stability.

While our results support a wider-reaching impact of bacteria on the stability of host proteins, it is worth noting that bacteria can selectively target components of the host proteostasis network. For example, *P. aeruginosa* produces toxins that target the mitochondrial unfolded protein response (UPR^MT^),^72^ a transcriptional pathway that ensures proper protein folding and clearance.^73^ As such, when the UPR^MT^ is overwhelmed or disrupted, proteotoxicity can occur, leading to accelerated protein aggregation.^74^ Our previous work demonstrated that *P. aeruginosa* and *E. coli* can also disrupt host proteostasis through the production of bacteria-derived protein aggregates.^16,23^ Additionally, *P. aeruginosa* FapC amyloids were shown to cross-seed with Aβ_1-42_.^75^ Interestingly, the *fap* operon is also present in the genomes of Burkholderiales, which include *Ralstonia* and *Achromobacter.*^76,77^ The *E. coli* amyloid, curli, enhanced the aggregation of α-synuclein *in vivo.*^78^ *K. pneumoniae* is also capable of producing amyloids,^79^ and was proteotoxic to all *C. elegans* lines (Figure 4). The underlying mechanisms of the bacteria-induced proteotoxicity remain elusive; however, it is likely that metastable proteins of bacterial origin sequester host chaperones. Such sequestration would diminish the capacity of the proteostasis network to buffer folding of endogenous proteins. A similar mechanism is supported by the organismal ability to buffer protein polymorphisms present within the host proteome, which is hindered by the introduction of misfolded proteins.^80^

A comprehensive understanding of individual bacterial residents of the human microbiota in the pathogenesis of neurodegenerative PCDs is crucial in developing effective interventions. Our study challenges the traditional approach of solely focusing on host-targeted therapeutics for neurodegenerative PCDs. Instead, modulating the gut microbiota may offer an effective strategy for preventing and managing these devastating diseases. Gut-targeted interventions will likely have to be implemented early in the disease or even prior to its onset. While our results indicate that bacteria generally affect host proteostasis, ultimately influencing the stability of host proteins, further studies are needed to decipher the role of proteoprotective and proteotoxic species in humans and devise approaches to control their levels.

## Acknowledgments

We thank Dr. Richard Morimoto and Sue Fox for providing *C. elegans* polyQ strains (Northwestern University), and Dr. Janine Kirstein (Leibniz Institute on Aging− Fritz Lipmann Institute) for providing *C. elegans* Aβ_1-42_ strain and the control. The other *C. elegans* strains were provided by the Caenorhabditis Genetics Center, which is funded by the NIH Office of Research Infrastructure Programs (P40 OD010440). Human bacterial isolates (listed in Table S1) were obtained through BEI Resources, NIAID, and NIH as part of the Human Microbiome Project. We thank Mark Gorelik for training RB and for assistance with computational analysis. We would like to thank the funders who have supported our work on the bacterial contribution to protein conformational diseases, namely the National Institute on Aging (R03AG070580), and the Infectious Diseases Society of America. Furthermore, we would like to thank the Department of Microbiology and Cell Science for start-up funding. Figure 1 was created using BioRender.com.

## Author contributions

DMC: conceived the project. ACW and DMC designed the study, analyzed data, and wrote the manuscript. ACW, RB, ASB performed the experiments. ACW and MB data visualization.

## STAR Methods

### Key resources table

*C. elegans* and bacterial strains that are not from the BEI repository can be found in Table 1.

### *C. elegans* maintenance and strains

*C. elegans* were maintained as previously described.^16^ For experiments that generated the data represented in the heat-map (Figure 2), nematodes were kept on *E. coli* OP50 for two days at 20°C, washed three times and transferred to indicated bacteria, where they were kept at 22.5°C for three days. For all other experiments, nematodes were plated on indicated bacteria as L1s at 22.5°C, where they remained until the time of assay, except for the experiment in Figure S3, in which worms were cultured in temperatures indicated in the figure. All *C. elegans* strains used in this study can be found in Table 1.

### Bacterial culture conditions

All bacteria were cultured at 37°C. Parenthesized numbers (1-5) in ST1 reflect the remaining conditions in which bacteria were grown in the heat-map (Figure 2): 1) anaerobically in reinforced clostridial broth (RCB) with Oxyrase, 2) facultative conditions (no shaking, tube filled to top) in brain heart infusion broth (BHI), 3) aerobic conditions, shaking at 220 revolutions per minute (RPM) in BHI, 4) anaerobic conditions on tryptic soy agar (TSA) supplemented with 5% defibrinated sheep’s blood, 5) aerobic conditions on TSA supplemented with 5% defibrinated sheep’s blood. *E. coli* OP50 controls were grown in a manner consistent with which experiment was performed. Bacteria were washed and resuspended in M9 before being seeded on NGM. All follow-up experiments were performed with bacteria grown on TSA supplemented with 5% defibrinated sheep’s blood at 37°C except for the intestinal confirmation experiment of the proteotoxic bacteria (Figure 4A, “Aggregates”). Except for *Prevotella* spp. which were grown under anaerobic conditions, bacteria were grown aerobically in follow-up experiments.

### Aggregate quantification

Fluorescent aggregates were quantified using Leica MZ10F Modular Stereo Microscope equipped with CoolLED pE300lite 365 dir mount STEREO with filter set ET GFP-MZ10F. Fluorescent aggregates in worms used to generate data for the heat-map (Figure 2, Table S1) were manually quantified after having been cultured as indicated in “*C. elegans* maintenance and strains”. For all other experiments, worms were plated on indicated bacteria as L1s and fluorescent aggregates were manually quantified after four days (intestinal polyQ44) or three days (muscle polyQ35, 40) of life as previously described.

### Motility assays

All motility assays were performed at room temperature as previously described.^16^ Time-off-pick (seconds) of transgenic nematodes harboring tissue-specific aggregating protein species was assessed as previously described,^16,17^ on three days (muscle polyQ, muscle Aβ_1-42_, TSΔ and respective controls) or four days (intestine polyQ, neuron polyQ, muscle α-synuclein and control) after having been plated on indicated bacteria as L1s.

### Live imaging

Nematodes expressing muscle-specific Aβ_1-42_ were plated as L1s on NGM containing *E. coli* OP50, *P. corporis* HM-1294, and *P. aeruginosa* PAO1 for three days were mounted, frozen to reduce background fluorescence, and imaged. Fluorescent and Nomarski images were taken using Zeiss Axio Observer 7, equipped as described previously.^16^ Images were processed as previously described.^16^

### Gel electrophoresis and western blotting

Worms were prepared for gel electrophoresis and western blotting which were performed as previously described with a few modifications.^16^ In brief, M9 was used to lift worms from NGM which were washed until superficial bacteria was removed. Worms were plated on unseeded NGM, were allowed to dry briefly and 50 worms were picked into 10 μL M9 with 1mM phenylmethylsulfonyl fluoride (PMSF) in non-skirted screw-cap microcentrifuge tubes and frozen overnight or longer at −80°C. Tubes were pulse-centrifuged and three, 1.5mm zirconium beads were added to each tube. The following two steps were repeated a minimum of two times; if large worm particulates were observed, these steps were repeated again with the entire sample set until no tubes contained worm pieces that were visible under a dissecting microscope: Samples were homogenized by placing tubes in a Bead Bug homogenizer at 280×10 rates per minute (RPM) for 90 seconds. Tubes were cooled on ice and centrifuged for 1-2 minutes on a low speed to re-settle worms and worm particulates. Samples were prepped for gel electrophoresis as previously described,^16^ and entire samples were loaded on 4-12% gradient sodium dodecylsulfate-polyacrylamide gels (SDS-PAGE) to separate proteins through electrophoresis. Proteins were transferred onto polyvinyl difluoride (PDVF) membrane, blocked with 5% nonfat milk powder in PBS-Tween-20 (PBST) and probed with Living Colors JL-8 monoclonal primary antibody (1:1000) for 48 hours at 4C on orbital shaker and underwent three, five-minute washes with 0.1% PBST followed by incubation 1:10,000 with goat-anti-mouse horse-radish peroxidase (HRP) conjugated secondary antibody for at least two hours followed by three, ten-minute washes with 0.1% PBST. Chemiluminescence was achieved by incubation with Clarity Western ECL substrate. Image J (v1.52) was used to analyze insoluble fractions.

### Quantification and statistical analysis

Quantification of aggregate counts per worm in Figure 2 were performed on a cohort of ten worms to corroborate the findings from the western blot analysis. In the rest of the figures, data are represented as the average number of aggregates or TOP (seconds) per worm obtained from two independent experiments for a total of 45-60 worms (Figure 3A “Aggregates”), 60 worms (Figure 4A “Aggregates”); three independent experiments for a total of 45 worms (Figure 3A “Motility”, Figure 3B, Figure 3C, Figure 4A “Motility”, Figure 4C “Aggregates”) or 30 worms (Figure 3D, Figure 3E, Figure 4B, Figure 4C “Motility”, Figure 4D, Figure 4E, Figure S1B, Figure S3); two independent experiments for a total of 20 worms (Figure S2). Data are described as statistically significant when p<0.05 as determined by Student’s t-test or one-way ANOVA followed by multiple comparison Dunnett’s post-hoc test performed using Graphpad Prism 8.4.3 or later. Degrees of significance are denoted by the use of asterisks such that *p<0.05, **p<0.01, ***p<0.001, ****p<0.0001. The heat map (Figure 1) is represented as data normalized to the control *E. coli* OP50. Error bars represent standard error of the mean.

### Heat map generation

Heatmap was constructed in R-studio using the ComplexHeatmap package from the Bioconductor project. The experimental data were fed into R-studio as an organized .CSV file. The heatmap is the graphical representation of this data, clustering the bacteria by phylum.

### Phylogenetic tree generation

Whole genome sequences were assembled using the Bacterial and Viral Bioinformatics Resources Center (BV-BCR) via Unicycler. All genomes were annotated in BV-BRC using Prokka. Phylogenetic analysis was conducted in BV-BRC based on concatenated alignments of 100 single-copy genes using mafft, and the results were assessed for maximum likelihood using RAxML. Sequences that did not have 100 complete single-copy genes due to poor sequencing quality were excluded.

## Key resources table

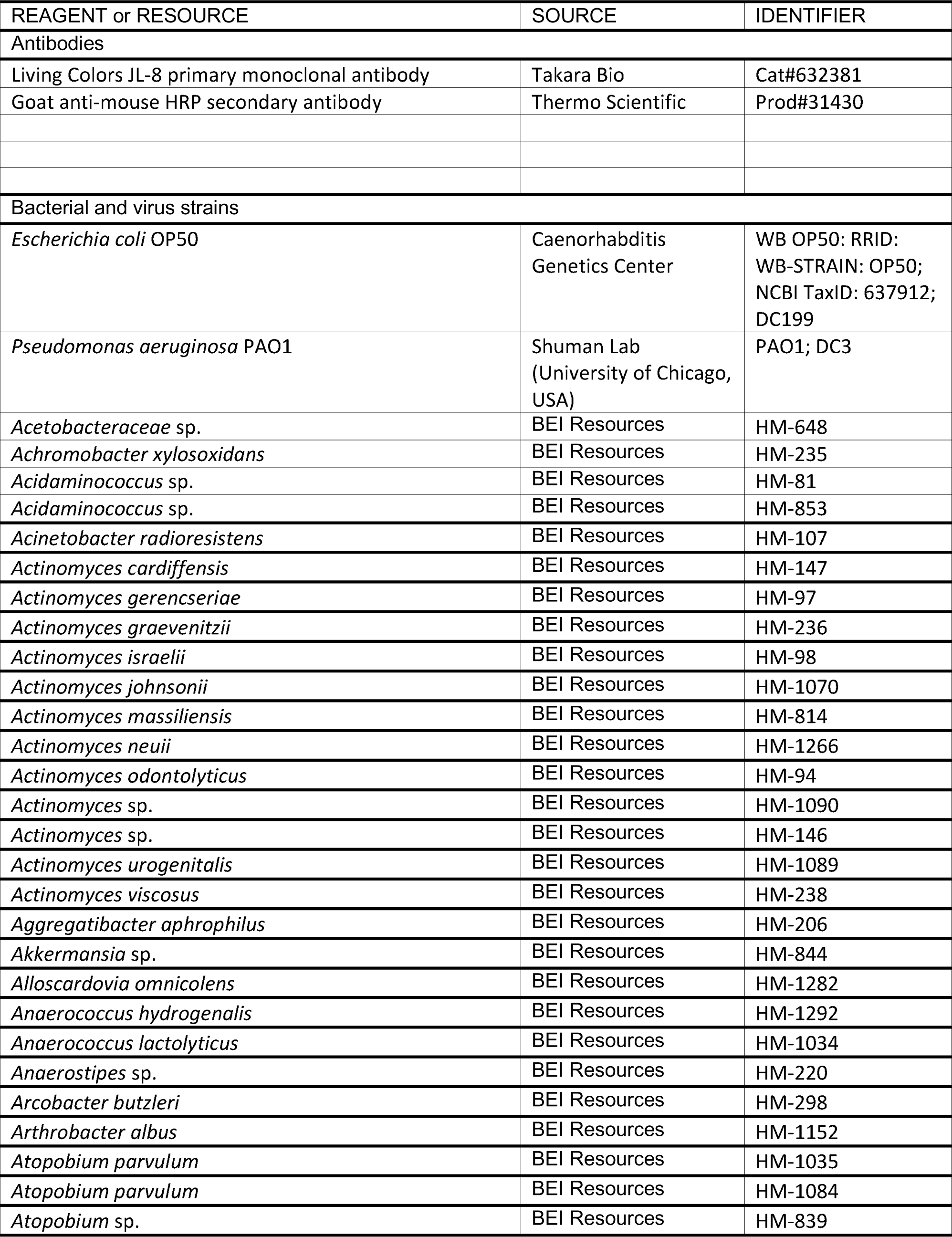
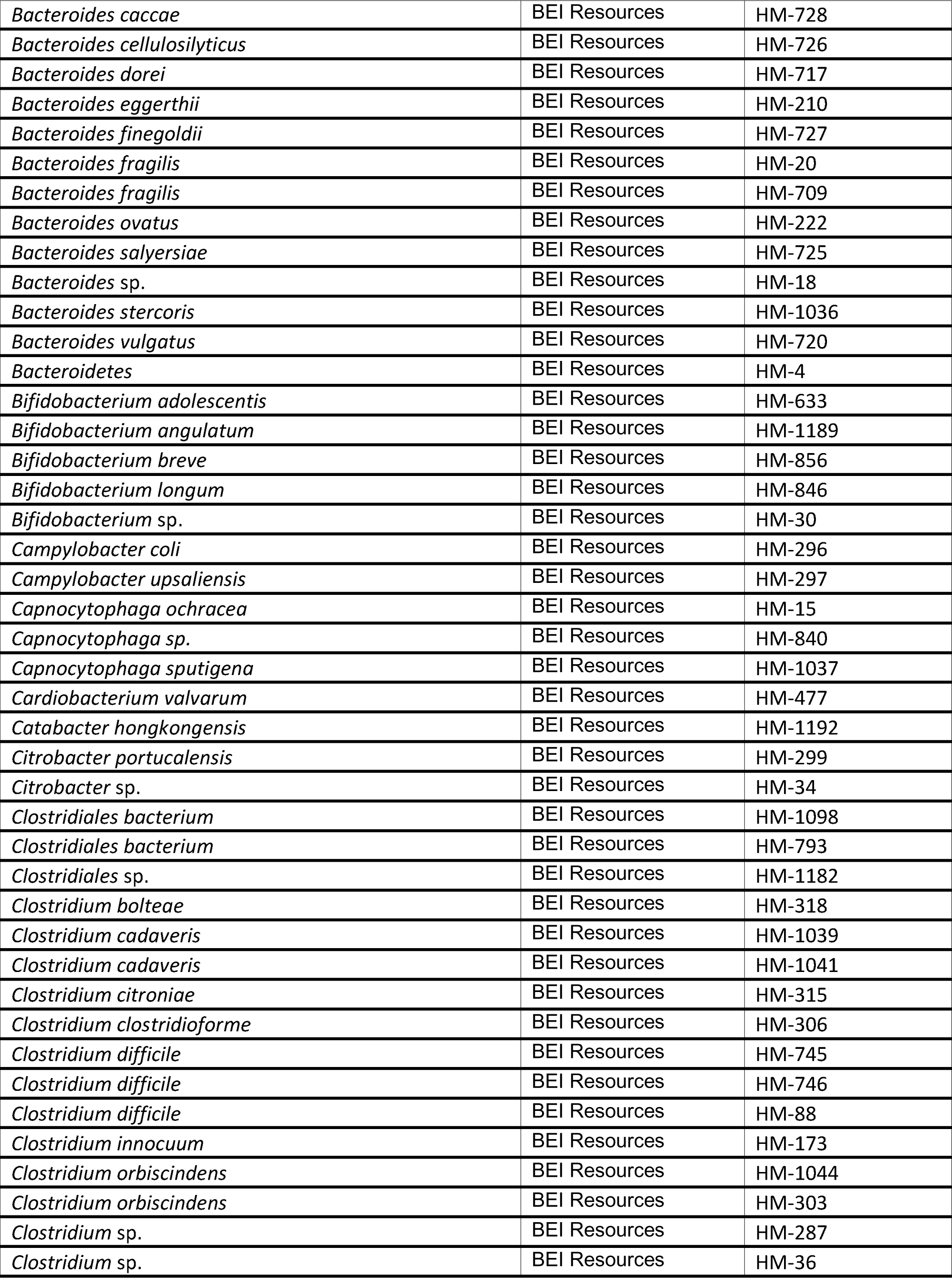
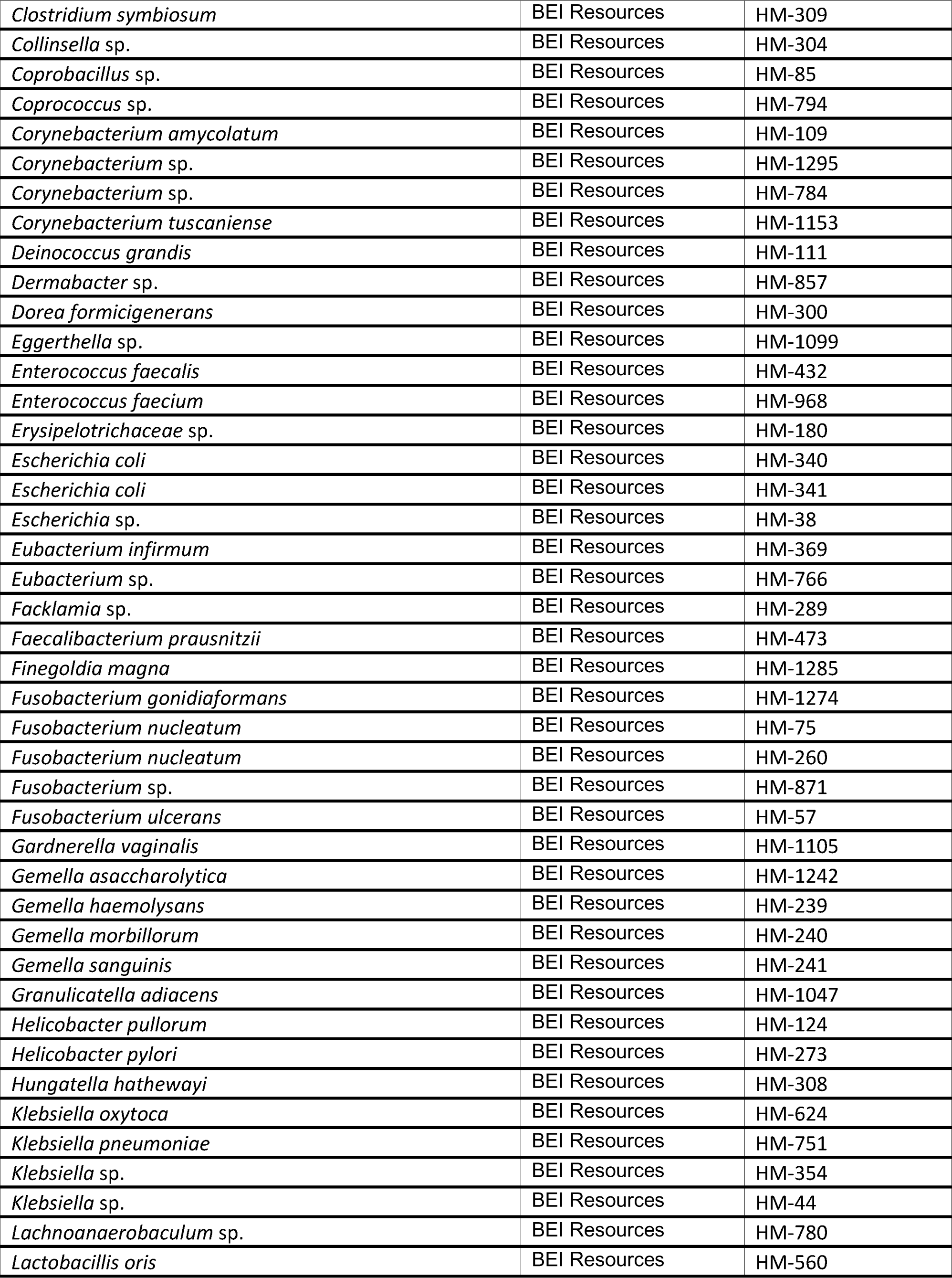
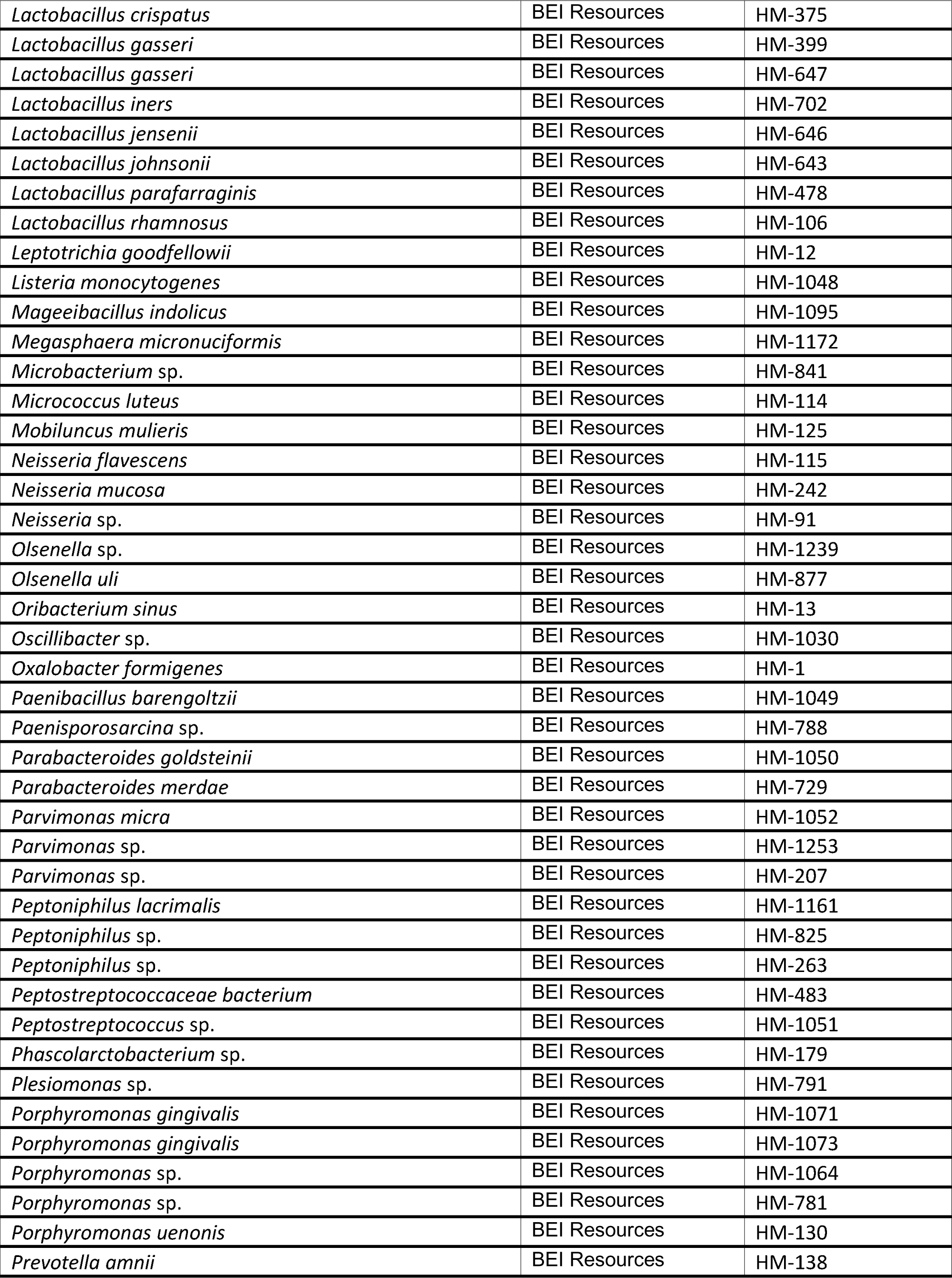
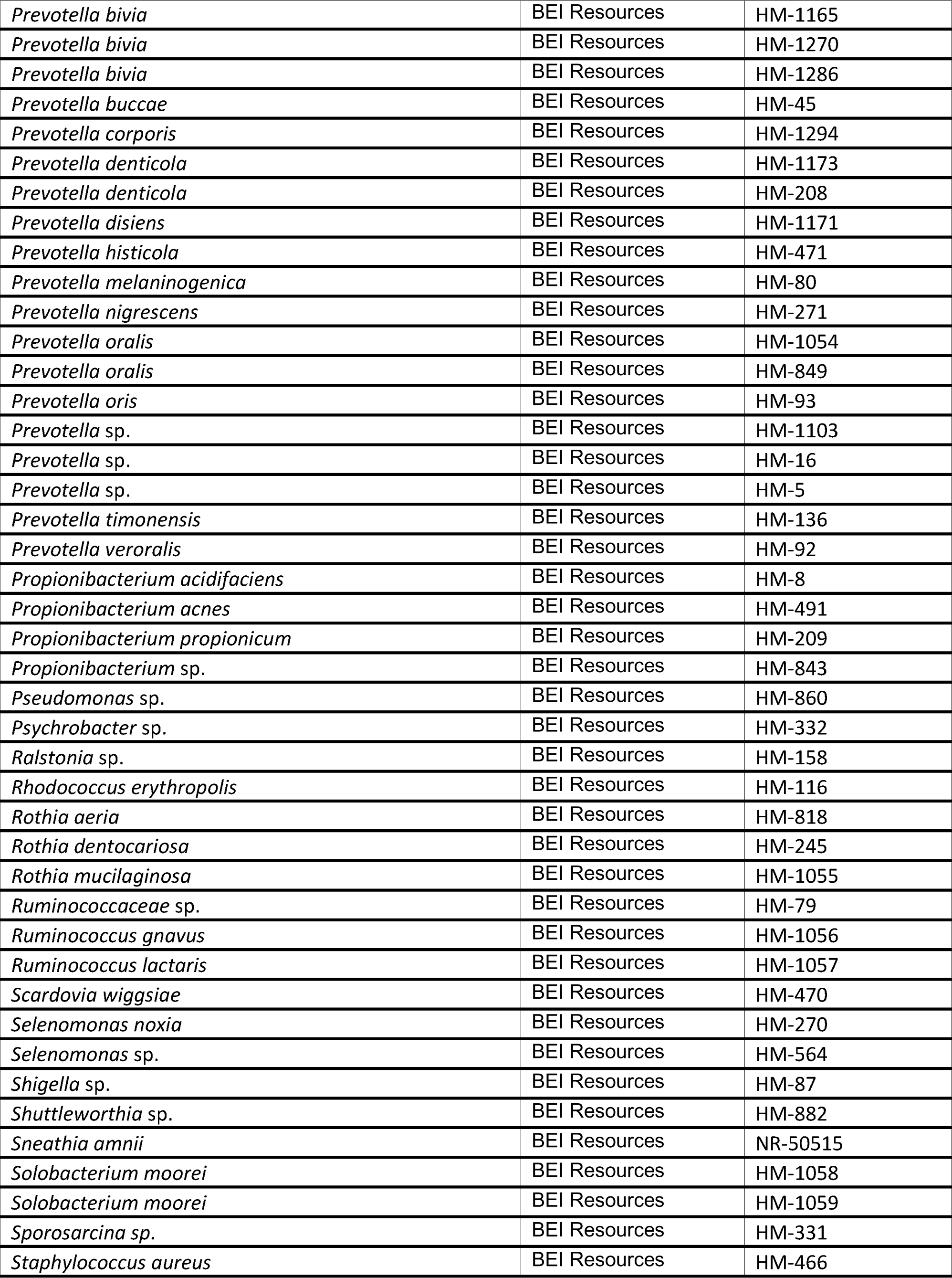
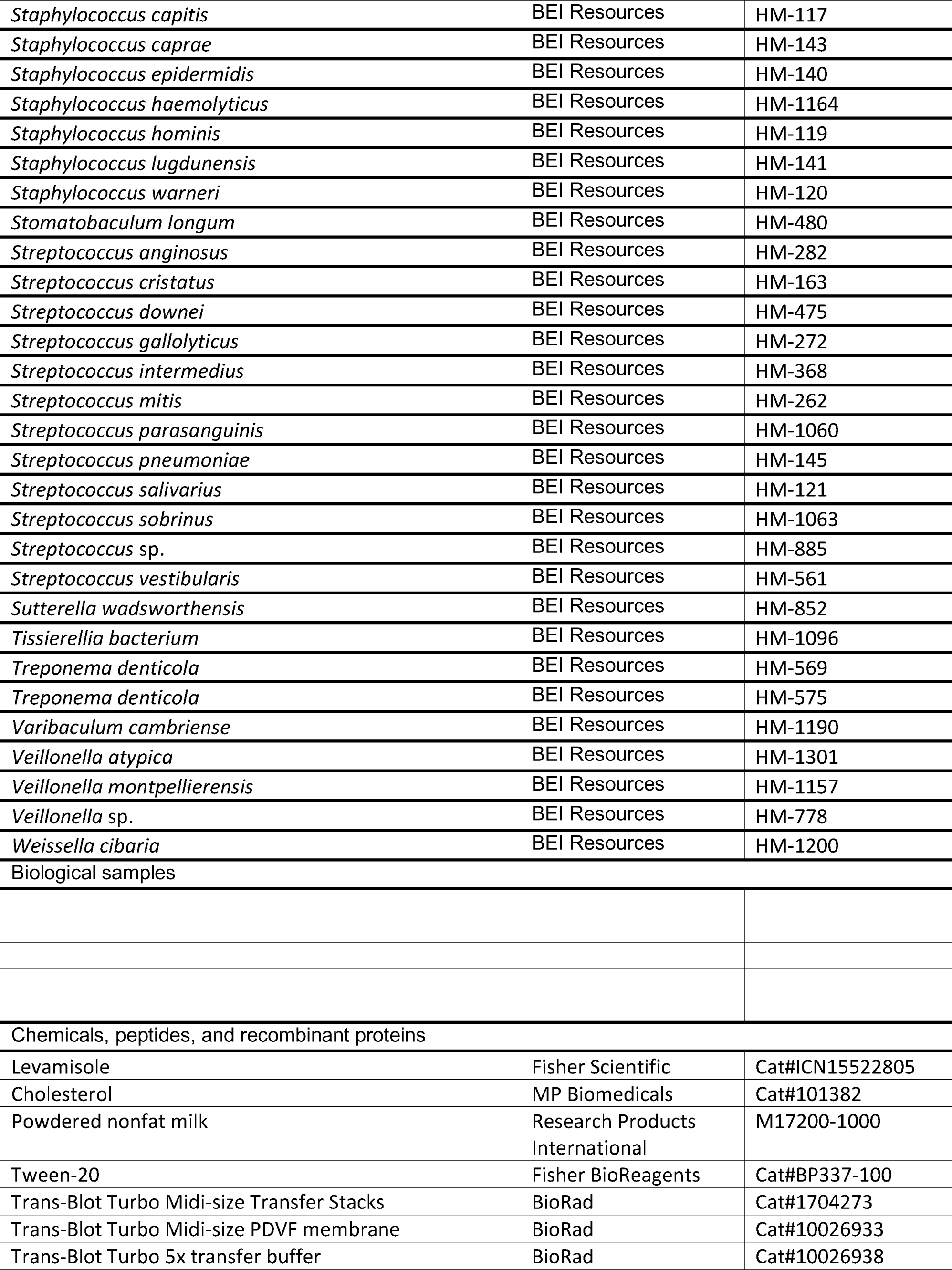
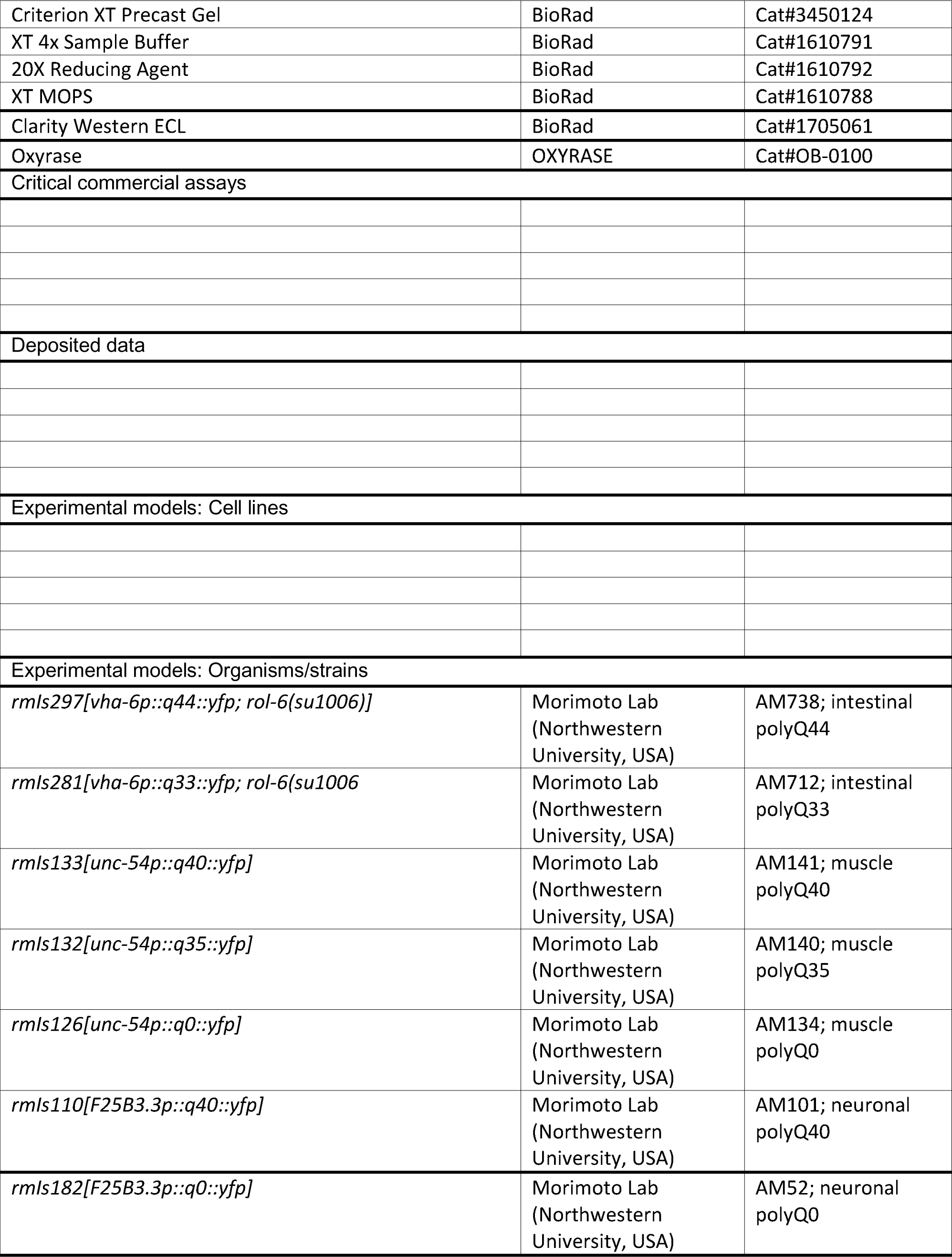
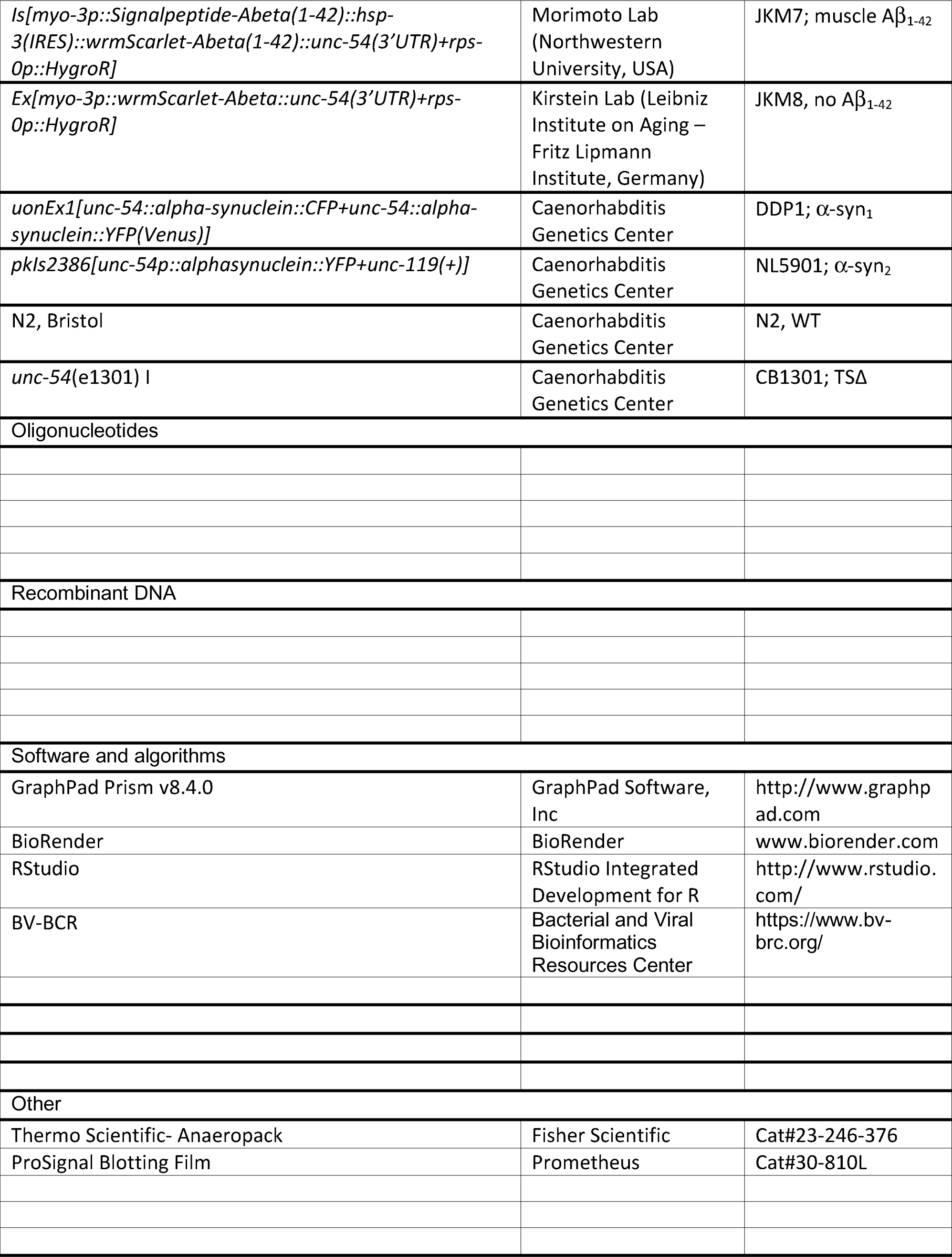

