## Supplementary material for "Identification of proteotoxic and proteoprotective bacteria that non-specifically affect proteins associated with neurodegenerative diseases": Table S1

| Bacteria | Strain | HM Number | Western | Count | Origin | Growth condition |
| --- | --- | --- | --- | --- | --- | --- |
| *Acetobacteraceae* sp. | AT-5844 | HM-648 | 1.59 | 1.89 | Infection | 3 |
| *Achromobacter xylosoxidans* | C54 | HM-235 | 2.76 | 2.70 | Respiratory Tract | 3 |
| *Acidaminococcus* sp. | D21 | HM-81 | 0.57 | 0.66 | GI Tract | 4 |
| *Acidaminococcus* sp. | HPA0509 | HM-853 | 0.10 | 0.47 | GI Tract | 1 |
| *Acinetobacter radioresistens* | SK82 | HM-107 | 0.72 | 0.59 | Skin | 3 |
| *Actinomyces cardiffensis* | F0333 | HM-147 | 2.27 | 1.60 | Oral Cavity | 2 |
| *Actinomyces gerencseriae* | F0344 | HM-97 | 1.12 | 0.97 | Oral Cavity | 2 |
| *Actinomyces graevenitzii* | C83 | HM-236 | 1.47 | 1.73 | Respiratory Tract | 2 |
| *Actinomyces israelii* | F0345 | HM-98 | 0.19 | 0.18 | Oral Cavity | 1 |
| *Actinomyces johnsonii* | F0510 | HM-1070 | 1.30 | 1.00 | Oral Cavity | 2 |
| *Actinomyces massiliensis* | F0489 | HM-814 | 4.60 | 2.07 | Oral Cavity | 2 |
| *Actinomyces neuii* | MJR8396A | HM-1266 | 2.27 | 2.13 | Vagina | 2 |
| *Actinomyces odontolyticus* | F0309 | HM-94 | 0.83 | 0.96 | Oral Cavity | 2 |
| *Actinomyces* sp. | S6-Spd3 | HM-1090 | 1.80 | 1.31 | Urogenital Tract | 1 |
| *Actinomyces* sp. | F0338 | HM-146 | 1.32 | 1.16 | Oral Cavity | 2 |
| *Actinomyces urogenitalis* | S6-C4 | HM-1089 | 0.59 | 0.65 | Vagina | 2 |
| *Actinomyces viscosus* | C505 | HM-238 | 1.64 | 1.86 | Respiratory Tract | 2 |
| *Aggregatibacter aphrophilus* | F0387 | HM-206 | 0.65 | 0.73 | Oral Cavity | 2 |
| *Akkermansia* sp. | KLE1605 | HM-844 | 0.39 | 0.53 | GI Tract | 1 |
| *Alloscardovia omnicolens* | CMW7705A | HM-1282 | 2.42 | 3.43 | Vagina | 1 |
| *Anaerococcus hydrogenalis* | MJR7738A | HM-1292 | 0.59 | 0.57 | Vagina | 1 |
| *Anaerococcus lactolyticus* | CC31C | HM-1034 | 0.17 | 0.53 | GI Tract | 1 |
| *Anaerostipes* sp. | 3_2_56FAA | HM-220 | 5.84 | 1.94 | GI Tract | 4 |
| *Arcobacter butzleri* | JV22 | HM-298 | 1.98 | 2.03 | GI Tract | 5 |
| *Arthrobacter albus* | DNF00011 | HM-1152 | 1.20 | 2.11 | Vagina | 3 |
| *Atopobium parvulum* | CC14Z | HM-1035 | 1.66 | 1.57 | GI Tract | 4 |
| *Atopobium parvulum* | DNF00906 | HM-1084 | 0.19 | 0.29 | Vagina | 4 |
| *Atopobium* sp. | F0494 | HM-839 | 2.54 | 1.74 | Oral Cavity | 1 |
| *Bacteroides caccae* | CL03T12C61 | HM-728 | 0.89 | 0.86 | GI Tract | 1 |
| *Bacteroides cellulosilyticus* | CL02T12C19 | HM-726 | 1.53 | 1.22 | GI Tract | 1 |
| *Bacteroides dorei* | CL02T00C15 | HM-717 | 28.27 | 4.35 | GI Tract | 1 |
| *Bacteroides eggerthii* | 1_2_48FAA | HM-210 | 0.97 | 1.03 | GI Tract | 1 |
| *Bacteroides finegoldii* | CL09T03C10 | HM-727 | 3.38 | 4.35 | GI Tract | 1 |
| *Bacteroides fragilis* | 3_1_12 | HM-20 | 63.50 | 4.35 | GI Tract | 1 |
| *Bacteroides fragilis* | CL07T00C01 | HM-709 | 61.59 | 4.35 | GI Tract | 1 |
| *Bacteroides ovatus* | 3_8_47FAA | HM-222 | 1.59 | 1.41 | GI Tract | 1 |
| *Bacteroides salyersiae* | CL02T12C01 | HM-725 | 0.49 | 0.80 | GI Tract | 1 |
| *Bacteroides* sp. | 2_1_22 | HM-18 | 0.34 | 0.58 | GI Tract | 1 |
| *Bacteroides stercoris* | CC31F | HM-1036 | 0.31 | 0.73 | GI Tract | 1 |
| *Bacteroides vulgatus* | CL09T03C04 | HM-720 | 0.49 | 0.79 | GI Tract | 1 |
| *Bacteroidetes* | F0058 | HM-4 | 0.19 | 0.38 | Oral Cavity | 4 |
| *Bifidobacterium adolescentis* | L2-32 | HM-633 | 0.88 | 1.28 | GI Tract | 1 |
| *Bifidobacterium angulatum* | F16_22 | HM-1189 | 0.73 | 0.72 | GI Tract | 1 |
| *Bifidobacterium breve* | HPH0326 | HM-856 | 2.81 | 3.84 | GI Tract | 4 |
| *Bifidobacterium longum* | 1-6B | HM-846 | 0.47 | 0.92 | GI Tract | 1 |
| *Bifidobacterium* sp. | 12_1_47BFAA | HM-30 | 1.20 | 2.17 | GI Tract | 4 |
| *Campylobacter coli* | JV20 | HM-296 | 0.68 | 0.95 | GI Tract | 2 |
| *Campylobacter upsaliensis* | JV21 | HM-297 | 0.35 | 0.95 | GI Tract | 2 |
| *Capnocytophaga ochracea* | F0287 | HM-15 | 0.30 | 0.46 | Oral Cavity | 2 |
| *Capnocytophaga sp.* | F0502 | HM-840 | 0.56 | 0.75 | Oral Cavity | 2 |
| *Capnocytophaga sputigena* | CC21_001D | HM-1037 | 1.60 | 1.26 | Unknown | 1 |
| *Cardiobacterium valvarum* | F0432 | HM-477 | 0.78 | 1.08 | Oral Cavity | 5 |
| *Catabacter hongkongensis* | AB8_9 | HM-1192 | 0.07 | 0.33 | GI Tract | 1 |
| *Citrobacter portucalensis* | 4_7_47CFAA | HM-299 | 3.09 | 3.74 | GI Tract | 3 |
| *Citrobacter* sp. | 30_2 | HM-34 | 8.17 | 6.45 | GI Tract | 3 |
| *Clostridiales bacterium* | S5-A14a | HM-1098 | 2.16 | 1.94 | Vagina | 4 |
| *Clostridiales bacterium* | HPP0074 | HM-793 | 2.94 | 3.10 | GI Tract | 1 |
| *Clostridiales* sp. | S9 PR-1 | HM-1182 | 1.48 | 1.19 | Vagina | 4 |
| *Clostridium bolteae* | WAL-14578 | HM-318 | 1.06 | 1.01 | GI Tract | 1 |
| *Clostridium cadaveris* | CC40_001C | HM-1039 | 2.12 | 3.02 | Unknown | 1 |
| *Clostridium cadaveris* | CC88A | HM-1041 | 0.65 | 0.84 | GI Tract | 1 |
| *Clostridium citroniae* | WAL-17108 | HM-315 | 0.48 | 0.74 | GI Tract | 1 |
| *Clostridium clostridioforme* | 2_1_49FAA | HM-306 | 0.36 | 0.48 | GI Tract | 1 |
| *Clostridium difficile* | 70-100-2010 | HM-745 | 5.06 | 3.98 | GI Tract | 1 |
| *Clostridium difficile* | 002-P50-2011 | HM-746 | 0.91 | 1.03 | Unknown | 1 |
| *Clostridium difficile* | NAP07 | HM-88 | 0.85 | 1.26 | GI Tract | 4 |
| *Clostridium innocuum* | 6_1_30 | HM-173 | 1.24 | 1.11 | GI Tract | 1 |
| *Clostridium orbiscindens* | CC43_001K | HM-1044 | 0.53 | 0.74 | Unknown | 1 |
| *Clostridium orbiscindens* | 1_3_50AFAA | HM-303 | 0.63 | 0.60 | GI Tract | 1 |
| *Clostridium* sp. | HGF2 | HM-287 | 2.41 | 1.68 | GI Tract | 1 |
| *Clostridium* sp. | 7_2_43FAA | HM-36 | 0.63 | 0.82 | GI Tract | 1 |
| *Clostridium symbiosum* | WAL-14163 | HM-309 | 2.31 | 1.67 | GI Tract | 1 |
| *Collinsella* sp. | 4_8_47FAA | HM-304 | 0.13 | 0.28 | GI Tract | 1 |
| *Coprobacillus* sp. | D6 (29_1) | HM-85 | 2.10 | 1.54 | GI Tract | 1 |
| *Coprococcus* sp. | HPP0048 | HM-794 | 0.33 | 0.65 | GI Tract | 1 |
| *Corynebacterium amycolatum* | SK46 | HM-109 | 1.21 | 1.41 | Skin | 5 |
| *Corynebacterium* sp. | CMW7794 | HM-1295 | 0.13 | 0.62 | Vagina | 3 |
| *Corynebacterium* sp. | HFH0082 | HM-784 | 0.27 | 0.61 | GI Tract | 3 |
| *Corynebacterium tuscaniense* | DNF00037 | HM-1153 | 0.67 | 1.43 | Vagina | 5 |
| *Deinococcus grandis* | SK125 | HM-111 | 1.62 | 1.46 | Skin | 5 |
| *Dermabacter* sp. | HFH0086 | HM-857 | 2.26 | 1.66 | GI Tract | 1 |
| *Dorea formicigenerans* | 4_6_53AFAA | HM-300 | 0.26 | 0.65 | GI Tract | 1 |
| *Eggerthella* sp. | MVA1 | HM-1099 | 0.97 | 1.01 | Vagina | 4 |
| *Enterococcus faecalis* | TX2137 | HM-432 | 2.16 | 3.18 | GI Tract | 2 |
| *Enterococcus faecium* | ERV102 | HM-968 | 0.08 | 0.57 | Oral Cavity | 2 |
| *Erysipelotrichaceae* sp. | 6_1_45 | HM-180 | 2.05 | 1.51 | GI Tract | 2 |
| *Escherichia coli* | MS 124-1 | HM-340 | 6.00 | 4.68 | GI Tract | 3 |
| *Escherichia coli* | MS 145-7 | HM-341 | 2.11 | 3.23 | GI Tract | 3 |
| *Escherichia* sp. | 3_2_53FAA | HM-38 | 2.02 | 3.39 | GI Tract | 3 |
| *Eubacterium infirmum* | F0142 | HM-369 | 0.17 | 0.63 | Oral Cavity | 1 |
| *Eubacterium* sp. | AS15 | HM-766 | 0.48 | 0.68 | Oral Cavity | 1 |
| *Facklamia* sp. | HFG4 | HM-289 | 1.58 | 1.41 | GI Tract | 4 |
| *Faecalibacterium prausnitzii* | KLE1255 | HM-473 | 2.14 | 1.76 | GI Tract | 1 |
| *Finegoldia magna* | GED7760A | HM-1285 | 1.34 | 1.02 | Vagina | 4 |
| *Fusobacterium gonidiaformans* | CMW8396 | HM-1274 | 2.34 | 1.59 | Vagina | 1 |
| *Fusobacterium nucleatum* | D11 | HM-75 | 0.70 | 0.90 | GI Tract | 1 |
| *Fusobacterium nucleatum* | F0401 | HM-260 | 0.08 | 0.45 | Oral Cavity | 1 |
| *Fusobacterium* sp. | CM21 | HM-871 | 1.36 | 1.08 | Oral Cavity | 4 |
| *Fusobacterium ulcerans* | 12_1B | HM-57 | 0.18 | 0.63 | GI Tract | 1 |
| *Gardnerella vaginalis* | JCP7275 | HM-1105 | 0.70 | 0.83 | Vagina | 5 |
| *Gemella asaccharolytica* | KA00071 | HM-1242 | 1.30 | 1.33 | Urogenital Tract | 1 |
| *Gemella haemolysans* | M341 | HM-239 | 0.90 | 0.64 | Respiratory Tract | 2 |
| *Gemella morbillorum* | M424 | HM-240 | 2.70 | 3.19 | Respiratory Tract | 2 |
| *Gemella sanguinis* | M325 | HM-241 | 2.03 | 2.27 | Respiratory Tract | 2 |
| *Granulicatella adiacens* | CC94D | HM-1047 | 2.00 | 1.53 | Unknown | 4 |
| *Helicobacter pullorum* | MIT 98-5489 | HM-124 | 0.30 | 0.69 | GI Tract | 2 |
| *Helicobacter pylori* | 83 | HM-273 | 0.24 | 0.60 | GI Tract | 2 |
| *Hungatella hathewayi* | WAL-18680 | HM-308 | 0.34 | 0.53 | GI Tract | 1 |
| *Klebsiella oxytoca* | MIT 10-5243 | HM-624 | 1.15 | 2.00 | Blood | 5 |
| *Klebsiella pneumoniae* | WGLW2 | HM-751 | 4.65 | 3.73 | Respiratory Tract | 3 |
| *Klebsiella* sp. | MS 92-3 | HM-354 | 2.53 | 3.69 | GI Tract | 3 |
| *Klebsiella* sp. | 1_1_55 | HM-44 | 2.85 | 3.32 | GI Tract | 3 |
| *Lachnoanaerobaculum* sp. | OBRC5-5 | HM-780 | 1.40 | 1.22 | Unknown | 1 |
| *Lactobacillis oris* | F0423 | HM-560 | 1.40 | 1.13 | Oral Cavity | 1 |
| *Lactobacillus crispatus* | EX849587VC07 | HM-375 | 1.27 | 1.02 | Vagina | 2 |
| *Lactobacillus gasseri* | EX336960VC02 | HM-399 | 1.15 | 1.02 | Vagina | 2 |
| *Lactobacillus gasseri* | 224-1 | HM-647 | 0.80 | 1.02 | Urogenital Tract | 1 |
| *Lactobacillus iners* | Lactin V 09 (V1)-c | HM-702 | 0.85 | 0.83 | Vagina | 1 |
| *Lactobacillus jensenii* | 208-1 | HM-646 | 1.12 | 1.00 | Vagina | 4 |
| *Lactobacillus johnsonii* | 135-1-CHN | HM-643 | 1.16 | 0.98 | Vagina | 1 |
| *Lactobacillus parafarraginis* | F0439 | HM-478 | 1.12 | 1.09 | Oral Cavity | 1 |
| *Lactobacillus rhamnosus* | LMS2-1 | HM-106 | 1.86 | 1.33 | GI Tract | 1 |
| *Leptotrichia goodfellowii* | F0264 | HM-12 | 0.21 | 0.78 | Oral Cavity | 1 |
| *Listeria monocytogenes* | CC70B | HM-1048 | 2.15 | 1.83 | GI Tract | 1 |
| *Mageeibacillus indolicus* | S7-24-11 | HM-1095 | 1.50 | 1.20 | Urogenital Tract | 4 |
| *Megasphaera micronuciformis* | DNF00954 | HM-1172 | 3.30 | 1.85 | Urogenital Tract | 4 |
| *Microbacterium* sp. | F0373 | HM-841 | 0.07 | 0.22 | Oral Cavity | 3 |
| *Micrococcus luteus* | SK58 | HM-114 | 0.86 | 0.93 | Skin | 3 |
| *Mobiluncus mulieris* | UPII 28-I | HM-125 | 0.23 | 0.60 | Vagina | 1 |
| *Neisseria flavescens* | SK114 | HM-115 | 1.90 | 2.22 | Skin | 3 |
| *Neisseria mucosa* | C102 | HM-242 | 0.35 | 0.51 | Respiratory Tract | 3 |
| *Neisseria* sp. | F0314 | HM-91 | 0.63 | 0.70 | Oral Cavity | 3 |
| *Olsenella* sp. | DNF00959 | HM-1239 | 1.36 | 1.09 | Vagina | 1 |
| *Olsenella uli* | MSTE5 | HM-877 | 1.75 | 1.31 | Oral Cavity | 1 |
| *Oribacterium sinus* | F0268 | HM-13 | 0.31 | 0.44 | Oral Cavity | 1 |
| *Oscillibacter* sp. | KLE 1728 | HM-1030 | 0.85 | 0.80 | GI Tract | 1 |
| *Oxalobacter formigenes* | OxCC13 | HM-1 | 0.21 | 0.45 | GI Tract | 1 |
| *Paenibacillus barengoltzii* | CC33_002B | HM-1049 | 0.15 | 0.55 | GI Tract | 3 |
| *Paenisporosarcina* sp. | HGH0030 | HM-788 | 0.47 | 0.97 | GI Tract | 5 |
| *Parabacteroides goldsteinii* | CC87F | HM-1050 | 6.77 | 4.29 | Unknown | 4 |
| *Parabacteroides merdae* | CL09T00C40 | HM-729 | 1.10 | 1.10 | GI Tract | 1 |
| *Parvimonas micra* | CC57A | HM-1052 | 0.54 | 0.70 | GI Tract | 4 |
| *Parvimonas* sp. | KA00067 | HM-1253 | 0.86 | 0.95 | Vagina | 4 |
| *Parvimonas* sp. | F0139 | HM-207 | 0.27 | 0.52 | Oral Cavity | 1 |
| *Peptoniphilus lacrimalis* | DNF00528 | HM-1161 | 0.21 | 0.54 | Vagina | 1 |
| *Peptoniphilus* sp. | BV3AC2 | HM-825 | 0.97 | 0.97 | Urogenital Tract | 1 |
| *Peptoniphilus* sp. | F0141 | HM-263 | 0.16 | 0.51 | Oral Cavity | 1 |
| *Peptostreptococcaceae bacterium* | ACC19a | HM-483 | 0.98 | 1.00 | Unknown | 1 |
| *Peptostreptococcus* sp. | CC14N | HM-1051 | 0.11 | 0.53 | Unknown | 4 |
| *Phascolarctobacterium* sp. | 3_1_syn4 | HM-179 | 1.25 | 1.68 | GI Tract | 1 |
| *Plesiomonas* sp. | HPP0020 | HM-791 | 0.31 | 0.83 | GI Tract | 1 |
| *Porphyromonas gingivalis* | F0568 | HM-1071 | 0.08 | 0.26 | Oral Cavity | 1 |
| *Porphyromonas gingivalis* | F0570 | HM-1073 | 0.36 | 0.57 | Oral Cavity | 1 |
| *Porphyromonas* sp. | W7784 | HM-1064 | 1.24 | 1.33 | Oral Cavity | 1 |
| *Porphyromonas* sp. | KLE1280 | HM-781 | 0.42 | 0.54 | Oral Cavity | 4 |
| *Porphyromonas uenonis* | UPII 60-3 | HM-130 | 0.90 | 0.83 | Urogenital Tract | 1 |
| *Prevotella amnii* | CRIS 21A-A | HM-138 | 0.77 | 0.63 | Urogenital Tract | 4 |
| *Prevotella bivia* | DNF00650 | HM-1165 | 0.93 | 0.92 | Urogenital Tract | 1 |
| *Prevotella bivia* | GED7880 | HM-1270 | 0.83 | 0.44 | Vagina | 1 |
| *Prevotella bivia* | GED7760C | HM-1286 | 0.49 | 0.70 | Urogenital Tract | 1 |
| *Prevotella buccae* | D17 | HM-45 | 0.87 | 0.61 | Oral Cavity | 1 |
| *Prevotella corporis* | MJR7716 | HM-1294 | 0.69 | 0.49 | Vagina | 1 |
| *Prevotella denticola* | DNF00960 | HM-1173 | 0.68 | 0.43 | Urogenital Tract | 1 |
| *Prevotella denticola* | F0289 | HM-208 | 0.10 | 0.54 | Oral Cavity | 1 |
| *Prevotella disiens* | DNF00882 | HM-1171 | 0.28 | 0.48 | Vagina | 1 |
| *Prevotella histicola* | F0411 | HM-471 | 0.64 | 0.57 | Oral Cavity | 1 |
| *Prevotella melaninogenica* | D18 | HM-80 | 0.51 | 0.60 | Oral Cavity | 1 |
| *Prevotella nigrescens* | F0103 | HM-271 | 1.33 | 1.44 | Oral Cavity | 1 |
| *Prevotella oralis* | CC98A | HM-1054 | 0.08 | 0.44 | Unknown | 1 |
| *Prevotella oralis* | HGA0225 | HM-849 | 0.09 | 0.41 | GI Tract | 1 |
| *Prevotella oris* | F0302 | HM-93 | 0.25 | 0.36 | Oral Cavity | 1 |
| *Prevotella* sp. | S7-1-8 | HM-1103 | 1.72 | 1.17 | Urogenital Tract | 1 |
| *Prevotella* sp. | F0295 | HM-16 | 0.68 | 0.59 | Oral Cavity | 1 |
| *Prevotella* sp. | F0108 | HM-5 | 0.81 | 0.72 | Oral Cavity | 1 |
| *Prevotella timonensis* | CRIS 5C-B1 | HM-136 | 1.20 | 1.01 | Urogenital Tract | 1 |
| *Prevotella veroralis* | F0319 | HM-92 | 0.31 | 0.56 | Oral Cavity | 4 |
| *Propionibacterium acidifaciens* | F0233 | HM-8 | 0.72 | 1.22 | Skin | 2 |
| *Propionibacterium acnes* | HL002PA3 | HM-491 | 2.07 | 1.60 | Skin | 2 |
| *Propionibacterium propionicum* | F0230 | HM-209 | 2.66 | 1.84 | Oral Cavity | 2 |
| *Propionibacterium* sp. | F0372 | HM-843 | 6.30 | 2.11 | Oral Cavity | 2 |
| *Pseudomonas* sp. | HPB0071 | HM-860 | 8.56 | 4.49 | GI Tract | 5 |
| *Psychrobacter* sp. | 1501 | HM-332 | 2.19 | 1.85 | Blood | 5 |
| *Ralstonia* sp. | 5_2_56FAA | HM-158 | 15.33 | 6.45 | GI Tract | 3 |
| *Rhodococcus erythropolis* | SK121 | HM-116 | 1.16 | 1.14 | Skin | 5 |
| *Rothia aeria* | F0474 | HM-818 | 0.75 | 0.73 | Oral Cavity | 2 |
| *Rothia dentocariosa* | M567 | HM-245 | 0.69 | 0.85 | Respiratory Tract | 2 |
| *Rothia mucilaginosa* | CC87LB | HM-1055 | 2.03 | 1.74 | GI Tract | 2 |
| *Ruminococcaceae* sp. | D16 | HM-79 | 2.19 | 1.76 | GI Tract | 1 |
| *Ruminococcus gnavus* | CC55_001C | HM-1056 | 0.88 | 0.85 | Unknown | 1 |
| *Ruminococcus lactaris* | CC59_002D | HM-1057 | 0.15 | 0.53 | Unknown | 1 |
| *Scardovia wiggsiae* | F0424 | HM-470 | 0.12 | 0.67 | Oral Cavity | 1 |
| *Selenomonas noxia* | F0398 | HM-270 | 0.64 | 0.68 | Oral Cavity | 1 |
| *Selenomonas* sp. | F0430 | HM-564 | 1.48 | 1.27 | Oral Cavity | 1 |
| *Shigella* sp. | D9 | HM-87 | 0.88 | 1.00 | GI Tract | 2 |
| *Shuttleworthia* sp. | MSX8B | HM-882 | 0.24 | 0.42 | Oral Cavity | 1 |
| *Sneathia amnii* | Sn35 | NR-50515 | 0.09 | 0.38 | Vagina | 1 |
| *Solobacterium moorei* | CC14Y | HM-1058 | 1.19 | 1.09 | Unknown | 1 |
| *Solobacterium moorei* | CC57E | HM-1059 | 0.60 | 0.90 | Unknown | 1 |
| *Sporosarcina sp.* | 2681 | HM-331 | 0.30 | 0.59 | Blood | 2 |
| *Staphylococcus aureus* | MRSA131 | HM-466 | 0.26 | 0.80 | Skin | 3 |
| *Staphylococcus capitis* | SK14 | HM-117 | 0.08 | 0.35 | Skin | 3 |
| *Staphylococcus caprae* | M23864:W1 | HM-143 | 0.19 | 0.48 | Skin | 3 |
| *Staphylococcus epidermidis* | BCM0060 | HM-140 | 0.24 | 0.36 | Skin | 3 |
| *Staphylococcus haemolyticus* | DNF00585 | HM-1164 | 1.69 | 1.53 | Vagina | 3 |
| *Staphylococcus hominis* | SK119 | HM-119 | 1.27 | 2.00 | Skin | 3 |
| *Staphylococcus lugdunensis* | M23590 | HM-141 | 0.31 | 0.64 | Skin | 3 |
| *Staphylococcus warneri* | SK66 | HM-120 | 0.27 | 0.41 | Skin | 3 |
| *Stomatobaculum longum* | ACC2 | HM-480 | 1.10 | 1.05 | Oral Cavity | 1 |
| *Streptococcus anginosus* | F0211 | HM-282 | 1.21 | 1.75 | Respiratory Tract | 5 |
| *Streptococcus cristatus* | F0329 | HM-163 | 1.40 | 2.18 | Oral Cavity | 3 |
| *Streptococcus downei* | F0415 | HM-475 | 3.71 | 3.90 | Oral Cavity | 3 |
| *Streptococcus gallolyticus* | TX20005 | HM-272 | 0.98 | 0.97 | Blood | 3 |
| *Streptococcus intermedius* | F0413 | HM-368 | 0.45 | 0.84 | Oral Cavity | 5 |
| *Streptococcus mitis* | F0392 | HM-262 | 0.15 | 0.52 | Oral Cavity | 3 |
| *Streptococcus parasanguinis* | CC87K | HM-1060 | 2.69 | 2.18 | GI Tract | 3 |
| *Streptococcus pneumoniae* | TCH843 | HM-145 | 1.28 | 1.04 | Respiratory Tract | 5 |
| *Streptococcus salivarius* | SK126 | HM-121 | 0.75 | 0.82 | Skin | 3 |
| *Streptococcus sobrinus* | W1703 | HM-1063 | 1.87 | 1.22 | Oral Cavity | 3 |
| *Streptococcus* sp. | CM7 | HM-885 | 0.23 | 0.50 | Oral Cavity | 3 |
| *Streptococcus vestibularis* | F0396 | HM-561 | 0.36 | 0.41 | Oral Cavity | 3 |
| *Sutterella wadsworthensis* | HGA0223 | HM-852 | 1.60 | 0.18 | GI Tract | 2 |
| *Tissierellia bacterium* | S5-A11 | HM-1096 | 2.34 | 1.52 | Vagina | 1 |
| *Treponema denticola* | SP37 | HM-569 | 5.10 | 2.25 | Oral Cavity | 4 |
| *Treponema denticola* | AL-2 | HM-575 | 3.65 | 3.76 | Oral Cavity | 1 |
| *Varibaculum cambriense* | AB12_3 | HM-1190 | 0.42 | 0.51 | GI Tract | 1 |
| *Veillonella atypica* | CMW7756B | HM-1301 | 0.27 | 0.44 | Vagina | 1 |
| *Veillonella montpellierensis* | DNF00314 | HM-1157 | 1.62 | 1.14 | Vagina | 1 |
| *Veillonella* sp. | ACP1 | HM-778 | 0.66 | 0.70 | Oral Cavity | 1 |
| *Weissella cibaria* | F16_1 | HM-1200 | 0.93 | 0.95 | GI Tract | 1 |
