## Supplementary figures and images for "Identification of proteotoxic and proteoprotective bacteria that non-specifically affect proteins associated with neurodegenerative diseases"

### Figure S1

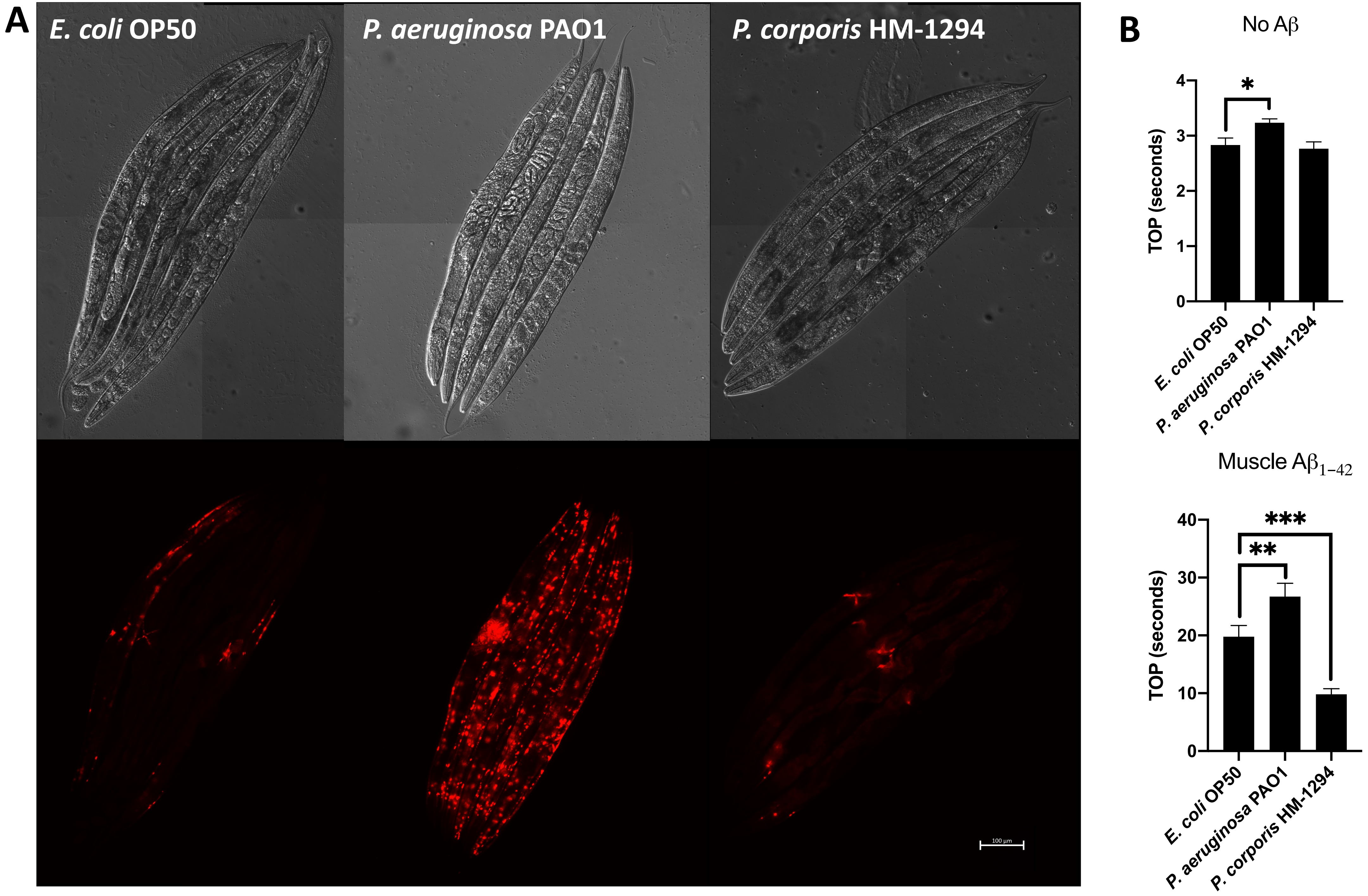

### Figure S2

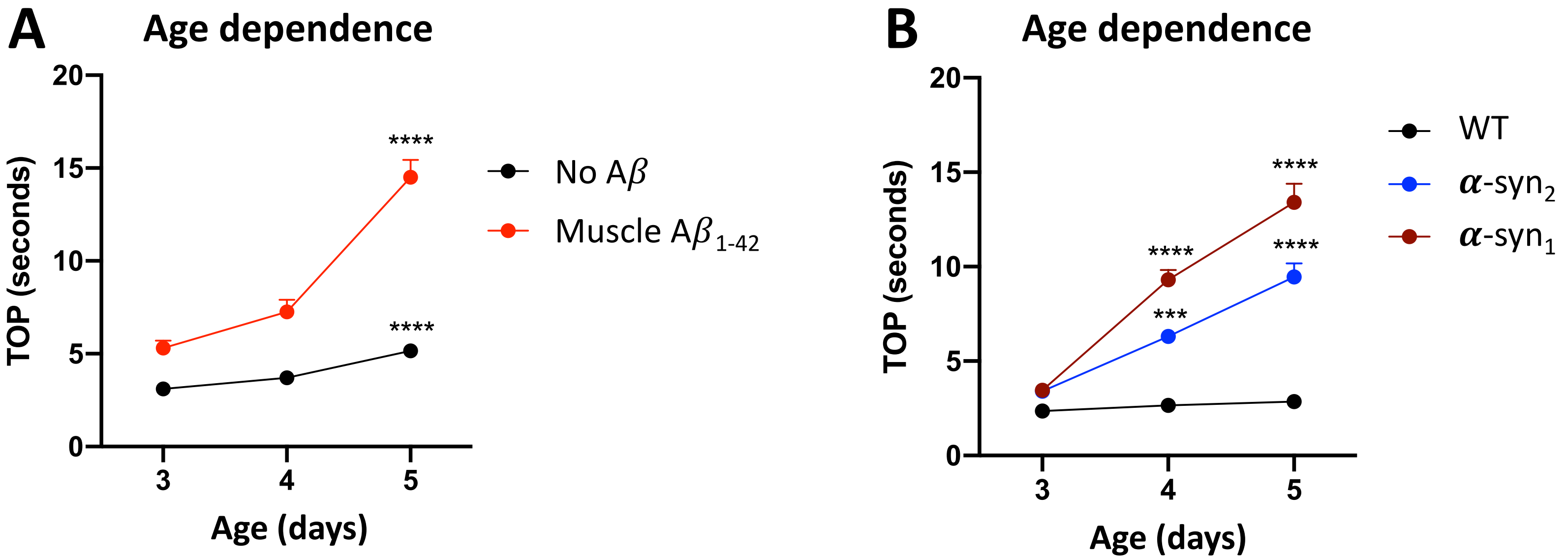

### Figure S3

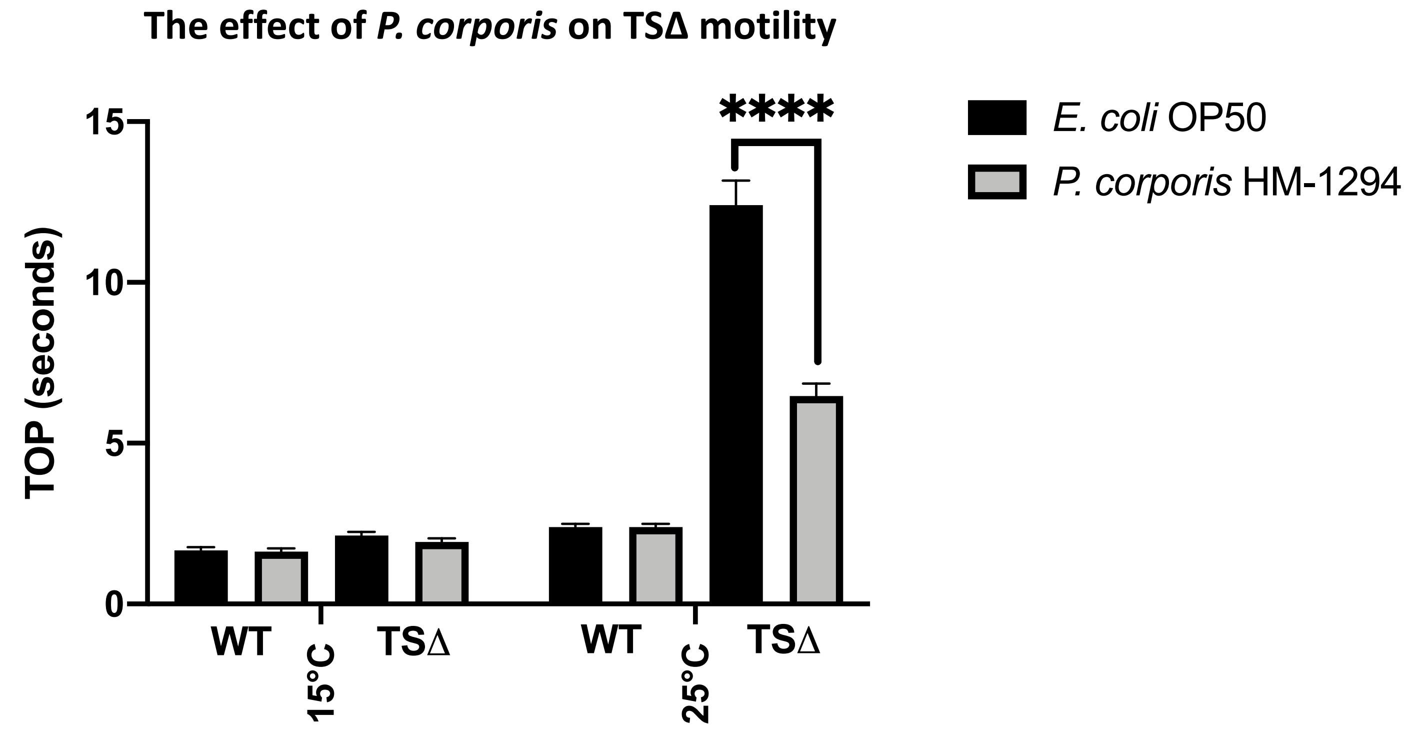

### Figure S4

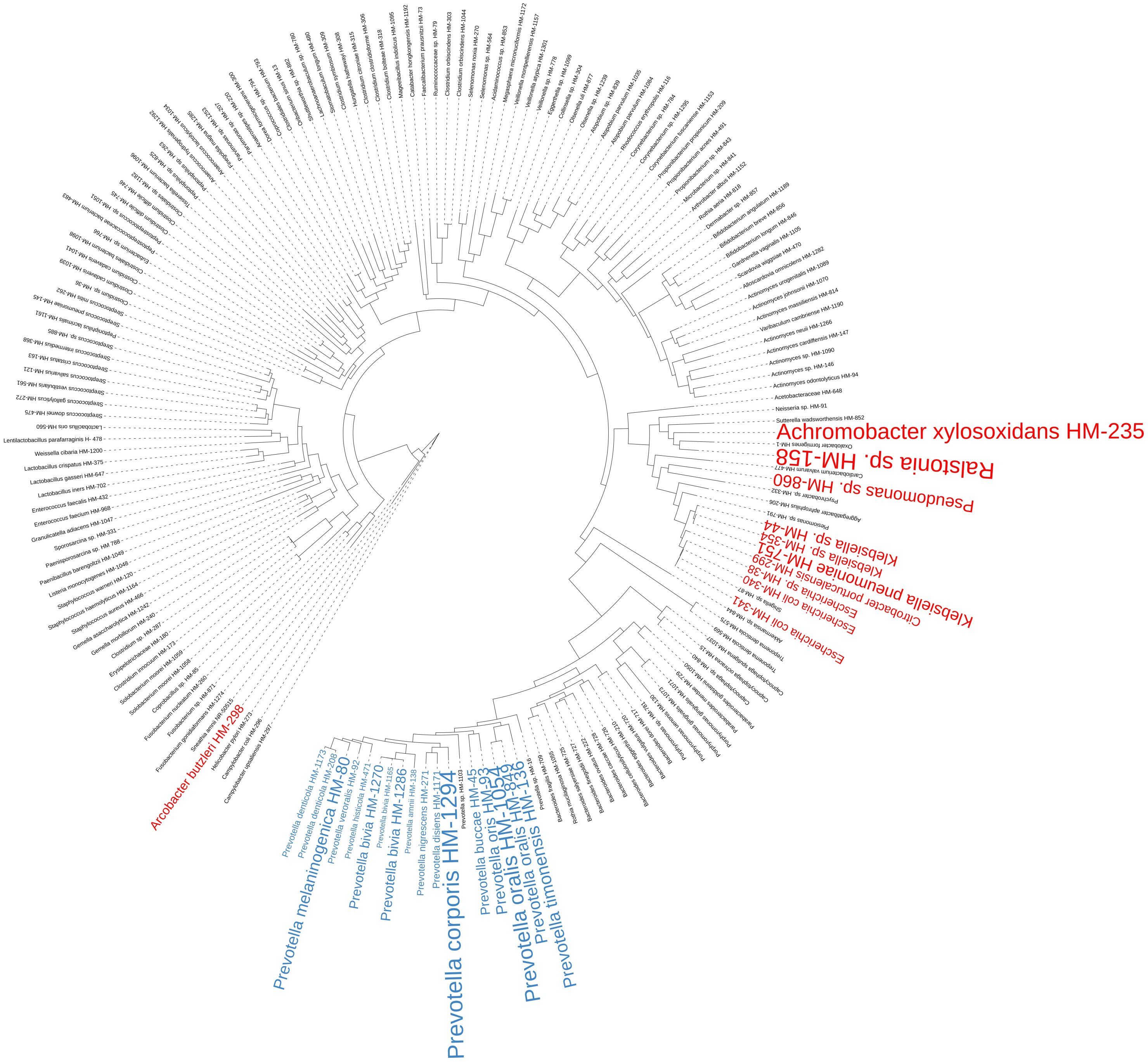
